# Reduced Non-Specific Binding of Super-Resolution DNA-PAINT Markers by Shielded DNA-PAINT Labeling Protocols

**DOI:** 10.1101/2024.06.03.597143

**Authors:** Evelina Lučinskaitė, Alexandre F. E. Bokhobza, Andrew Stannard, Anna Meletiou, Chris Estell, Steven West, Lorenzo Di Michele, Christian Soeller, Alexander H. Clowsley

## Abstract

The DNA-based single molecule super-resolution imaging approach, DNA-PAINT, can achieve nanometer resolution of single targets. However, the approach can suffer from significant non-specific background signals originating from non-specifically bound DNA-conjugated DNA-PAINT secondary antibodies as shown here. Using dye-modified oligonucleotides the location of DNA-PAINT secondary antibody probes can easily be observed with widefield imaging prior to beginning a super-resolution measurement. This reveals that a substantial proportion of DNA probes can accumulate, non-specifically, within the nucleus, as well as across the cytoplasm, of cells. Here, Shielded DNA-PAINT labeling is introduced, a method using partially or fully double-stranded docking strand sequences, prior to labeling, in buffers with increased ionic strength to greatly reduce non-specific interactions in the nucleus as well as the cytoplasm. This new labeling approach is evaluated against various conditions and it is shown that applying Shielded DNA-PAINT can reduce non-specific events ∼5 fold within the nucleus. This marked reduction in non-specific binding of probes during the labeling procedure is comparable to results obtained with unnatural left-handed DNA albeit at a fraction of the cost. Shielded DNA-PAINT is a straightforward adaption of current DNA-PAINT protocols and enables nanometer precision imaging of nuclear targets with low non-specific background.

## Background

Super-resolution microscopy is a powerful technique to achieve beyond diffraction limited imaging of biological targets using light alone^1–3^. DNA-PAINT belongs to a branch of super-resolution imaging called single molecule localization microscopy (SMLM) and has been shown to yield single nanometer localization precision under appropriate conditions^4–6^. The approach circumvents limitations of alternative SMLM approaches, namely photobleaching and the requirement for poorly controllable photoswitchable dyes and buffers. Instead, in DNA-PAINT, markers are conjugated with short oligonucleotides called docking strands. Complementary sequences with a dye modified end, called imagers, are introduced into the imaging buffer. These imagers are free to stochastically, and importantly reversibly, bind to available docking site/s on the docking strand. It is the transient immobilization of the imager to these sites that we detect as a bright single molecule event via a camera. The duration of these binding events can be fine-tuned by altering the number of matching base pairs^7–9^ or adjusting the ionic strength of the buffer^10^. DNA binding kinetics and sequence design are well understood which has led to the design of highly versatile nanobiotechnological tools^11–13^. Despite these advancements there are still limitations, including the possibility of non-specific binding interactions that may occur to varying levels in DNA-PAINT imaging^7^. This is a serious concern when attempting to image proteins with unknown/variable/sparse distributions in distinguishing real from spurious binding events as it may not be possible to identify and remove such events post-hoc by computational discrimination. The problem of non-specific events is greatly exacerbated and highly noticeable when imaging targets within and around the nucleus of cells. Improved methods for super-resolution detection of nuclear factors will enable new analyses of cellular events. Several nuclear processes (*e*.*g*. the synthesis and processing of RNA) take place in defined membrane-less compartments^14,15^. The composition of these compartments is unclear, including the arrangement and stoichiometry of their components. Such information is critical for accessing mechanistic details. Methods like DNA-PAINT are potentially powerful for addressing such questions but have historically suffered from high nuclear background signals, limiting their application. Recently, an approach using left-handed DNA (L-DNA) was put forward as a potential solution to reduce these non-specific interactions^16^. L-DNA does not hybridize to natural right-handed DNA (R-DNA) which should reduce non-specific interactions resulting from imagers or docking strands hybridizing to endogenous DNA in the nucleus. The commercial procurement of L-DNA is significantly more expensive per nucleotide than conventional R-DNA. This makes purchasing longer sequences, which are often required for more advanced super-resolution approaches, research development and experimentation, prohibitive. In this study, we introduce a new labeling procedure, called Shielded DNA-PAINT, that greatly reduces the levels of non-specific interactions within cells when preparing samples for DNA-PAINT imaging using conventional and easy to procure right-handed DNA. Shielded DNA-PAINT involves partially, or fully, double stranding DNA-PAINT antibodies in higher salt containing solutions prior to applying them to biological samples leading to a reduction in the levels of non-specific uptake caused by those markers. We additionally present the capability to shield short 11 nt DNA-PAINT antibodies using an excess of complementary strands that, due to their size, can be simply washed off prior to imaging.

## Results

### Retention of DNA-PAINT Markers in the Nucleus of Cells

Immunohistochemistry labeling using commercially available primary and secondary antibodies should ideally exhibit negligible non-specific interactions around the cytoplasm or nucleus of biological samples. Figure 1a, which shows the mitochondria located signal from anti-Tom20 antibodies where mitochondria are arranged around the location of the (here invisible) nucleus, demonstrates the expected fluorescent labeling outcome. For DNA-PAINT experiments our laboratory has generally adopted dye modified oligonucleotides as docking strands in order to ascertain labeling by widefield imaging prior to often time-consuming super-resolution imaging^17,18^. Working with COS-7 cells we noticed a strong nuclear fluorescence signal when we replaced a normal commercial Cy3 secondary antibody with one of our Cy3 dye modified DNA-PAINT secondaries, Figure 1b. This nuclear background signal was also clearly visible when omitting the primary antibody, Figure 1c. By fully hybridizing the docking strand during the labeling procedure the nuclear signal arising from non-specifically bound secondary antibodies (ABs) was greatly reduced, Figure 1d. However, this fully double-stranded configuration precludes DNA-PAINT super-resolution imaging as the docking site is not accessible to imager strands. Prompted by these observations we investigated the sources of DNA-PAINT backgrounds quantitatively.

**Figure 1.**
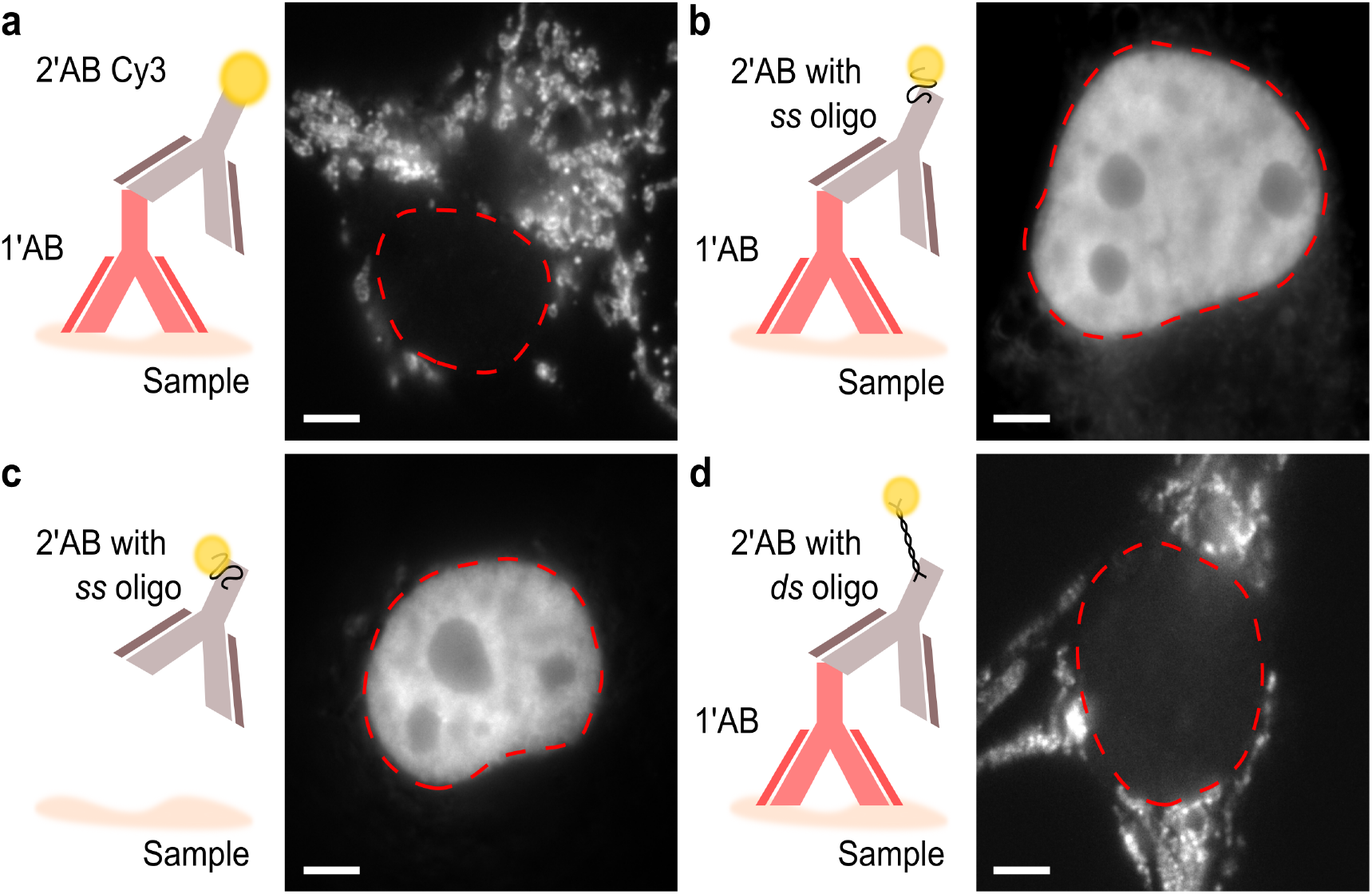
Various labeling approaches applied to COS-7 cells. **a**. A primary monoclonal antibody targeting mitochondrial target Tom20 and a commercial Cy3 secondary antibody labels only the target. Note that the nuclear region is clear from fluorescence. **b**. The commercial secondary antibody is replaced with a Cy3 dye modified DNA-PAINT oligonucleotide conjugated to a blank secondary antibody. This results in a visibly strong non-specific nuclear signal. **c**. ALempting to label the cells with the DNA-PAINT secondary antibody only shows the same level of non-specific uptake within the nucleus. **d**. A fully double-stranded DNA-PAINT secondary significantly reduces non-specific accumulation of markers within the nucleus. Red dashed lines indicate the nuclear perimeter. Scale bar: 5 μm. Images (b-d) taken with antibodies conjugated to docking strand D2 (see Table 1).

### Imager-only and Marker-based DNA-PAINT Background Signals

Previous work on nuclear DNA-PAINT backgrounds focused on backgrounds resulting from imagers non-specifically binding in the nucleus^16^. We replicated such observations by quantitatively investigating the non-specific nuclear binding of different imager sequences in fixed COS-7 cells free from any docking site containing antibody markers, Figure 2a. A nominal 0.1 nM imager concentration was applied to the sample and imaged under DNA-PAINT super-resolution imaging conditions, see also Supplementary Table 1. In these experiments the imager binding rate is specified as detected events per μm^-2^ per 1000 camera frames, with each frame lasting 100 ms, which we call 1 *edu* (event density unit, i.e. 1 *edu* = 0.01 events μm^-2^ s^-1^). This takes into account that in DNA-PAINT the number of events is proportional to the area sampled and the duration of imaging as photo-bleaching is effectively eliminated (as long as “docking strand site loss” is small). Photodamage has previously been observed to result in docking strand site loss in DNA-PAINT imaging of DNA-origami^19^. In a primary/secondary antibody system, where there are many markers, the loss is small at typical excitation powers and relatively short (<1 hour) acquisition time as indicated by a near constant event rate for the duration of an experiment. All event densities are given as the mean ± standard error of the mean.

**Figure 2.**
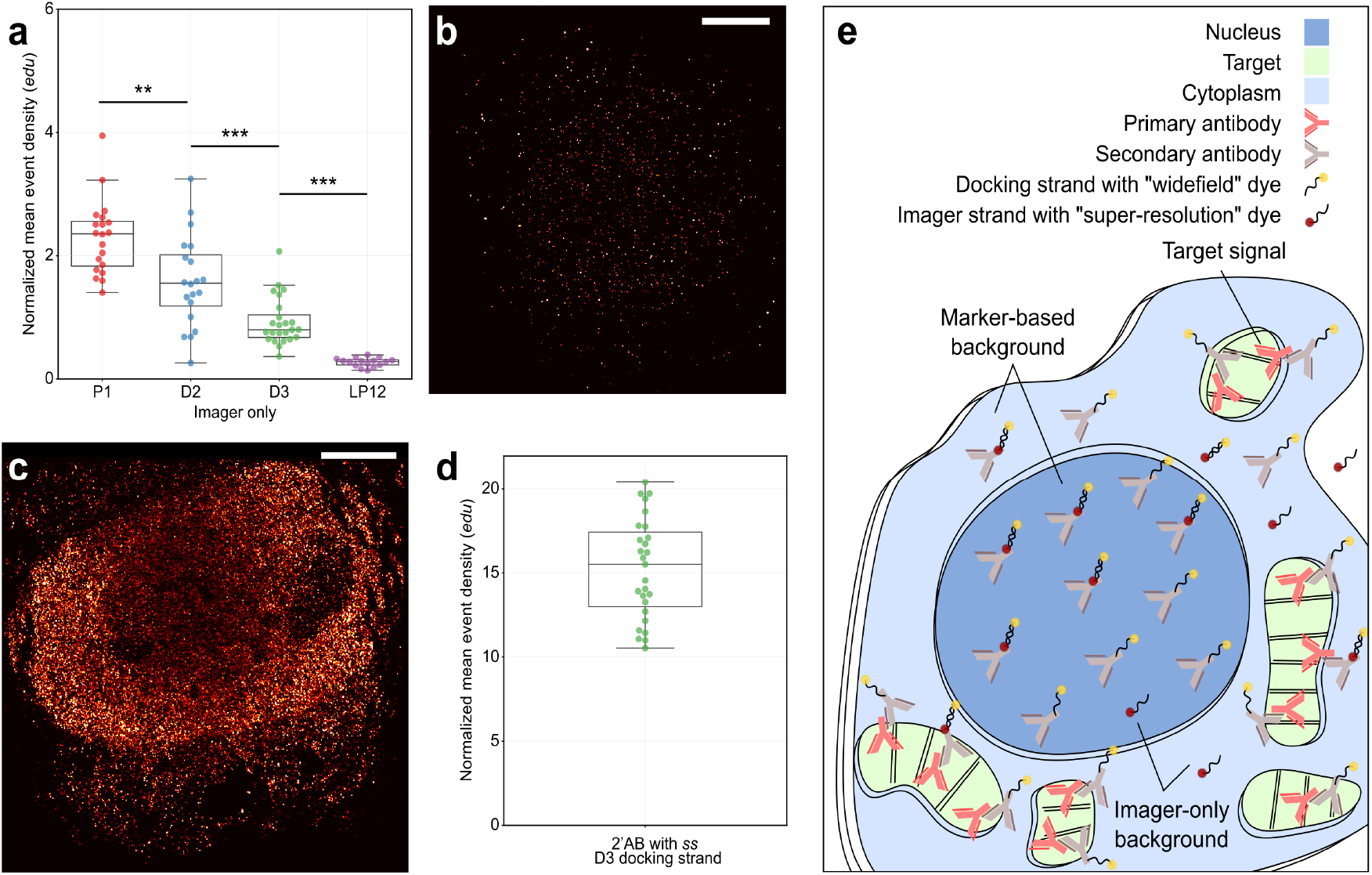
Pathways leading to non-specific localization event detection of oligonucleotides. **a**. A plot of the normalized mean event density (*edu*) for non-specific localizations measured across nuclear regions of unlabeled cells for various imagers. Three right-handed oligonucleotide imager sequences P1 imager, D2 imager and D3 imager are shown and have decreasing nuclear presence (2.30 ± 0.13, 1.58 ± 0.17, 0.93 ± 0.08 *edu* respectively). The LP12 imager, a left-handed non-natural oligonucleotide, has the least number of detections within the nucleus, 0.27 ± 0.02 *edu, (*20, 20, 24 & 16 cells, *n* = 3). **b**. An example rendered image of a nuclear region in an unlabeled cell exposed to 0.1 nM D3 imager for 5 k frames. **c**. A COS-7 cell incubated with a D3 DNA-PAINT conjugated secondary antibody (2’AB) only, imaged and rendered as in *b*. **d**. A plot of normalized mean event localization density for cells labeled non-specifically in this manner show an increase in detected events by over an order of magnitude, 15.3 ± 0.6 *edu* (27 cells, *n* = 3), when compared to corresponding D3 imager-only experiments. **e**. Schematic showing possible routes of single molecule event detections in biological samples. Marker-based background is caused by the non-specific attachment of conjugated oligonucleotide ABs during labeling. A much smaller, imager-only, background is caused by the non-specific, stochastic transient trapping of imagers on or inside the cell. All values given as mean ± standard error of the mean. Boxplot points represent the mean value obtained per measured cell, and line indicates the median. Independent two-tailed t-tests indicate the following levels of significance: ** p ≤ 0.01 and *** p ≤ 0.001. Scale bars: 5 μm.

The often used P1 imager (10 nt), first used in the original introduction of the DNA-PAINT methodology^5^, exhibited slightly elevated nuclear event densities, 2.30 ± 0.13 *edu* in comparison to D2 imagers (10 nt), 1.58 ± 0.17 *edu*. The shorter D3 imager (9 nt) sequence had fewer non-specific events at 0.93 ± 0.08 *edu*. Recently, non-natural left-handed DNA sequences (L-DNA) have been shown to facilitate the imaging of nuclear targets^16^. We therefore also tested a L-DNA sequence reported to behave favorably, called the LP12 imager (9 nt)^16^. As expected, this strand exhibited the lowest levels of non-specific binding within the nucleus, having an event density of 0.27 ± 0.02 *edu*, see Table 1 for the full list of sequences. Figure 2b shows a rendered example of localization data for a D3 imager acquisition from a nuclear region. Sporadic and sparse event detections can be seen.

**Table 1.**
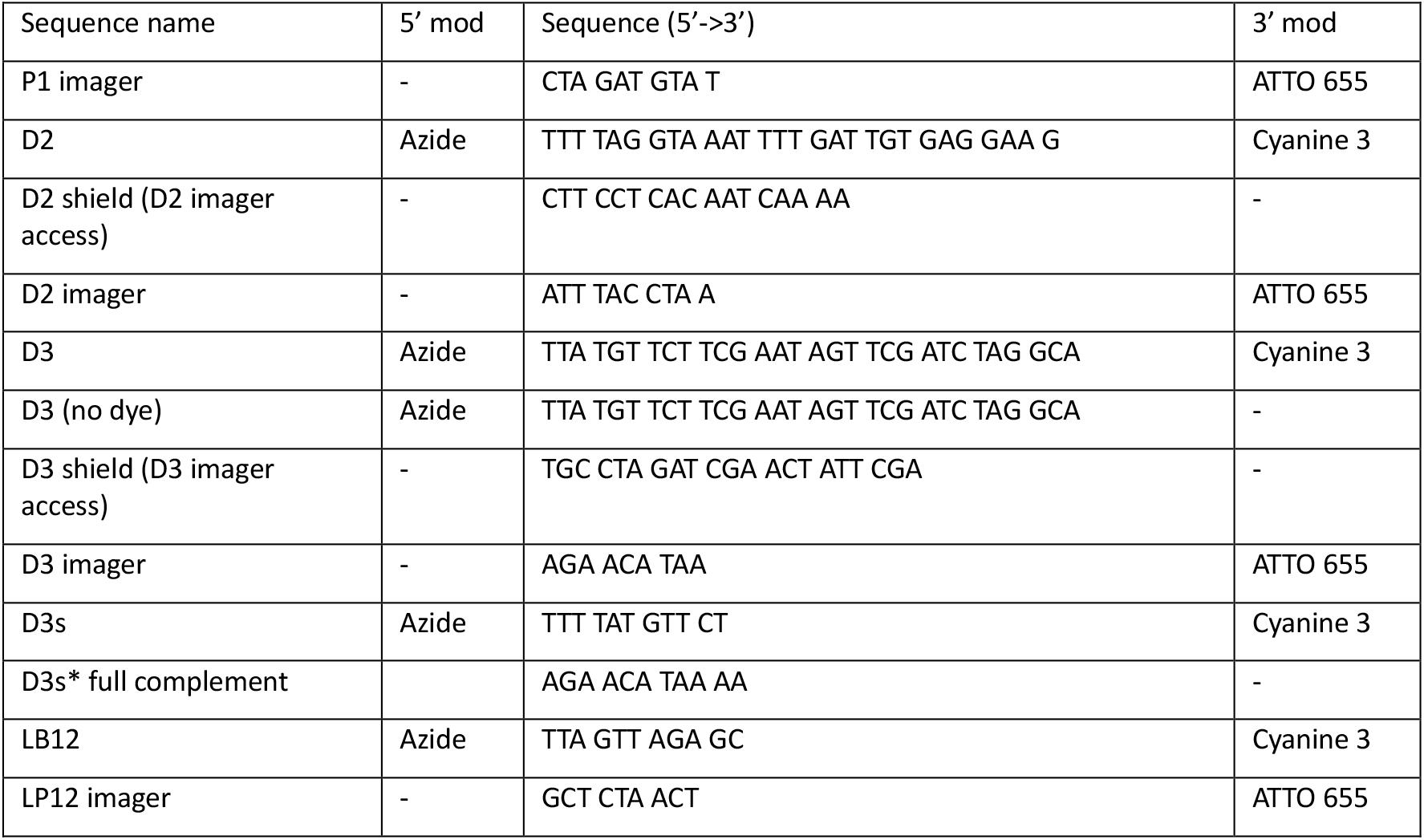
Oligonucleotide sequences and their modifications (mod).

By contrast, application of a DNA-PAINT secondary antibody conjugated with D3 docking strands exhibits a much larger number of events across an otherwise unlabeled cell, when probed with D3 imager, see Figure 2c. Experiments with a D3 docking strand conjugated secondary AB only (no primary ABs) yielded a nuclear event density of 15.3 ± 0.6 *edu*, Figure 2d, an ∼16 fold increase in nuclear event density compared to using D3 imagers only in cells not incubated with DNA-PAINT antibodies. These data suggest that the non-specific binding of DNA-PAINT antibodies in the sample (compare also Figure. 1c) leads to more than an order of magnitude higher background event densities than the direct binding of imagers to nuclear constituents. In other words, DNA-PAINT marker-associated backgrounds in practice far dominate over the “imager-only” backgrounds that were previously studied^16^. We therefore investigated DNA-PAINT marker-associated backgrounds more systematically and sought methods to reduce it.

Figure 2e schematically highlights the different origins of specific versus non-specific event localizations in DNA-PAINT super-resolution imaging. Conceptually, recorded events arise from three types of interactions, (i) imagers binding non-specifically to endogenous constituents of the cell, (ii) imagers binding (as designed) to complementary docking strands on DNA-PAINT antibodies that themselves are non-specifically bound in the cell and (iii) imagers binding to docking strands on DNA-PAINT antibodies bound to their intended target, i.e. primary antibodies of the targeted species. We call the background arising from non-specifically bound DNA-PAINT antibodies the “marker-based background” versus the “imager-only background” and finally, the desired signal corresponding to the presence of the primary ABs we term the “target signal”. In experiments with just imagers (no primary and secondary ABs) we measure solely imager-only background. In experiments with no primaries but using DNA-PAINT secondaries, we measure the sum of marker-based and imager-only backgrounds. In experiments with primary ABs and secondary DNA-PAINT ABs we measure the sum of all 3 signal contributions, but their contributions vary in different regions of the cell. Data interpretation is greatly simplified by one of the signals dominating over the others. For example, in cell regions where abundant target is concentrated (here mitochondria) the target signal dominates over the backgrounds as we show below.

### Shielded DNA-PAINT Labeling

To reduce the comparatively prominent nuclear marker-based background contributions identified in our experiments we modified the incubation step in which secondary antibodies are applied during the labeling procedure. We hypothesized that since a fully double-stranded sequence greatly reduced the uptake of DNA-PAINT markers into the nucleus (Figure 1d) then a partially double-stranded (*δds*) system may similarly reduce marker-based background. Importantly, a *δds* docking strand leaves access for an imager to bind to the single-stranded (*ss*) unshielded portion of the sequence, Figure 3a, making DNA-PAINT feasible.

**Figure 3.**
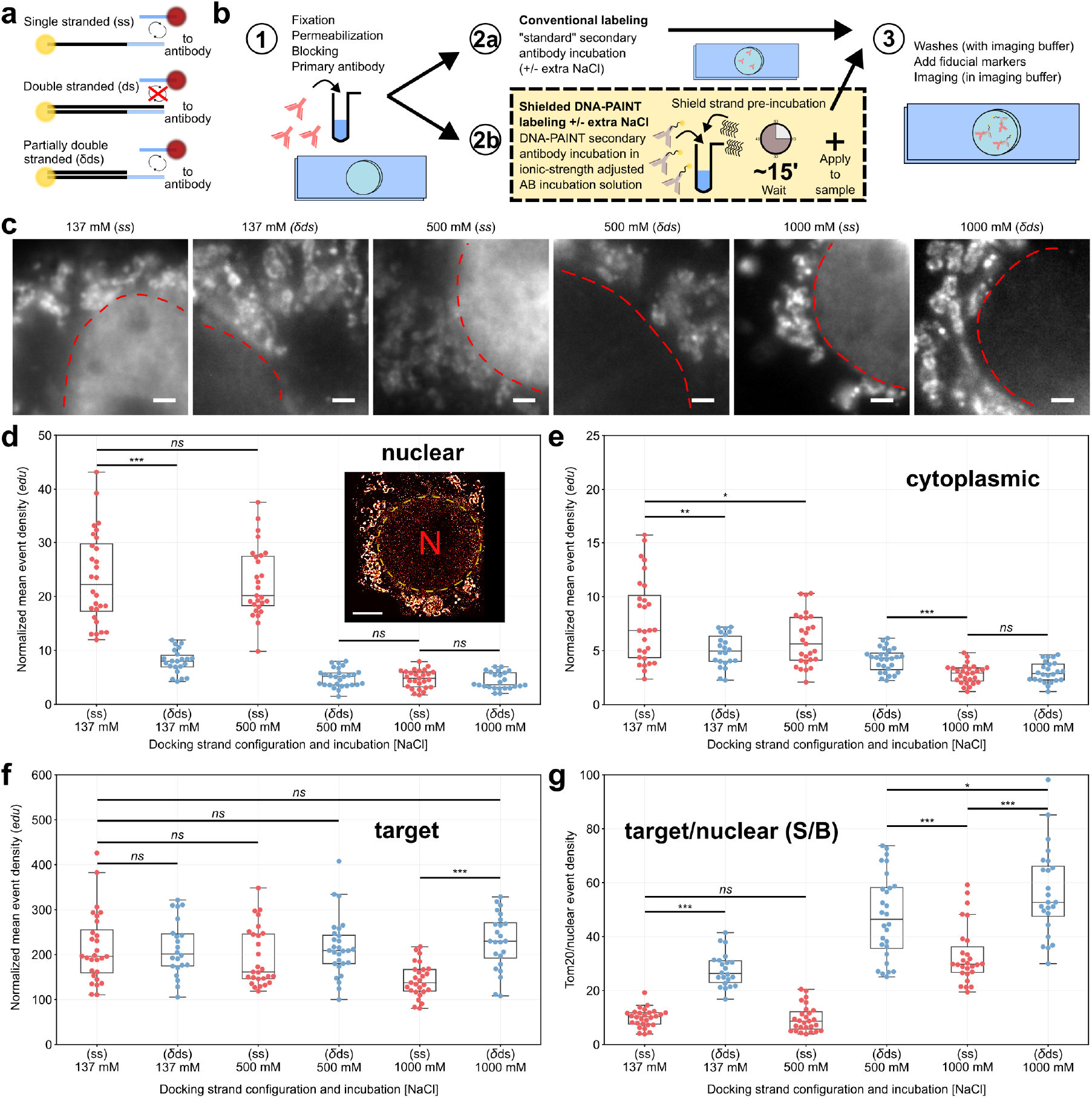
Comparison between conventional secondary antibody labeling and Shielded DNA-PAINT labeling using adapted incubation solutions for reduced non-specific interactions. **a**. DNA-PAINT imagers can freely hybridize to docking sites on single-stranded (*ss*) oligonucleotides at target locations. When the strand at the target location is double-stranded (*ds*), imagers can no longer bind. In a partially double-stranded (*δds*) system DNA-PAINT super-resolution measurements are still possible. **b**. Workflow for labeling biological samples. (1) the sample is fixed, permeabilized, blocked and labeled with a primary antibody. (2) The sample is then either labeled with a DNA-PAINT secondary in (a) a normal antibody incubation solution (PBS based) typically containing 137 mM NaCl or incubated in (b) a modified solution containing up to 1M NaCl with complementary ‘shield’ strands that form a partially double-stranded (*δds*) system aimed at reducing the non-specific binding of the secondary antibody during labeling. (3) The sample is washed in imaging buffer, fiducials are added and the sample finally imaged with appropriate imager sequences. **c**. Widefield images of COS-7 cells labeled for Tom20 using Cy3 dye modified D3 DNA-PAINT secondary with the various incubation solutions containing, from left to right: *ss* and *δds* at increasing NaCl concentrations (137, 500, 1000 mM). Normalized mean event densities calculated from localization events detected for the different labeling conditions in **d**. nuclear regions & **e**. cytoplasmic regions show decreased levels of non-specific binding with all *δds* systems and with 1000 mM NaCl. Target event densities of Tom20, **f**, showed similar levels in *ss* and *δds* configurations. A slight reduction in *ss* state at 1 M was observed. **g**. The ratio of nuclear event densities to target showed improved signal to background (S/B) for *δds* compared to *ss*, (28, 22, 27, 28, 27 & 24 cells, *n* = 3). Boxplot points represent the mean value obtained per measured cell, and line indicates the median. Independent two-tailed t-tests indicate the following levels of significance: *ns* p > 0.05, * p ≤ 0.05, ** p ≤ 0.01 and *** p ≤ 0.001. Scale bars: c) 2 μm, inset in d) 5 μm.

With this motivation the labeling procedure was adapted in a process we term Shielded DNA-PAINT labeling. In this technique, we pre-incubate the DNA-PAINT secondary ABs with excess ‘*shield’* strands purposely designed to create a *δds* system, Figure 3b, prior to adding the shielded secondary ABs in this incubation solution to the sample. In addition, we investigated the use of higher ionic strength incubation solutions, prompted by the use of higher ionic strength to reduce non-specific backgrounds when using DNA-barcoded antibodies^20^. As argued there, increasing ionic strength should help reduce electrostatic interactions between DNA and charged constituents of the cell. In addition, higher ionic strength will also stabilize the hybridization of the shield strands with the DNA-PAINT AB docking strands to form the shielded *δds* docking strand configuration. To investigate the combined effects of shielding and ionic strength, we evaluated both *ss* and *δds* DNA systems at three NaCl concentrations: 137 (as in a standard PBS solution), 500 and 1000 mM. In the *ss* configuration (i.e. omitting the shield strands) each of the widefield images exhibited a clear nuclear signal indicative of non-specific uptake of the DNA-PAINT secondary antibody, Figure 3c. When the DNA-PAINT secondary was incubated using the Shielded DNA-PAINT approach, i.e. with docking strands in a *δds* state, the signal intensity from the nucleus in widefield images was greatly reduced already at the standard NaCl concentration of 137 mM. At higher salt concentrations (500-1000 mM) the nuclear region appeared virtually free from obvious signal with Shielded DNA-PAINT.

To confirm these observations quantitatively in super-resolution data, event densities of DNA-PAINT localization data were analyzed for each incubation condition using COS-7 cells labeled with an anti-Tom20 primary antibody and a Cy3-modified D3 docking strand conjugated secondary antibody to show Tom20 distribution in mitochondria.

Cells labeled in this way were probed with a D3 imager that binds in the *ss* domain of the D3 docking strand, see also Supplementary Figure 1. The dissociation rate, *k*_*OFF*_, was similar between *ss* and *δds* configurations indicating the presence of the ‘shield’ strand had minimal effect on imager binding times, Supplementary Figure 2a. To quantify the background and signal components, event densities were evaluated in 3 different regions of the cell, (1) in the nuclear region, (2) in the mitochondrial target regions where Tom20 is known to be highly expressed and (3) in cytoplasmic regions distal to mitochondria to quantify cytoplasmic background (for further details of the analysis see Methods and Supplementary Figure 3, and for super-resolution examples see Supplementary Figure 4). Event densities in the nuclear region are expected to be dominated by marker-based background, in the target regions (mitochondria) would be dominated by target signal whereas in cytoplasmic regions away from mitochondria marker-based cytoplasmic backgrounds should dominate.

Cells labeled “conventionally” in a PBS based antibody incubation solution containing standard 137 mM NaCl and unshielded *ss* D3 docking strand ABs exhibited the highest levels of non-specific signals within the nucleus with an event density of 23.4 ± 1.6 *edu*, Figure 3d. Application of Shielded DNA-PAINT in the same buffer saw a significant decrease to 7.9 ± 0.5 *edu* (p << 0.001). There was no significant difference when incubating *ss* D3 at 500 mM NaCl compared to *ss* at 137 mM NaCl (p = 0.483). Nuclear event densities were lowest in *δds* systems containing 500 or 1000 mM NaCl and interestingly also in *ss* at 1000 mM NaCl, 4.9 ± 0.3, 4.3 ± 0.3 and 4.7 ± 0.3 *edu*, respectively (p << 0.001 compared to *δds* 137 mM NaCl). A smaller ∼2.6x reduction was observed in cytoplasmic background regions, Figure 3e, when comparing *ss* at 137 mM to *δds* 1 M NaCl event densities (p << 0.001).

The target event densities measured within mitochondria and resulting therefore mostly from “target signal” (compare also Fig. 2e) remained similar across the various labeling approaches at ∼215 *edu*, Figure 3f. A general reduction in target event densities were however observed in *ss* systems where the NaCl concentration was increased from 137 mM, 215.4 ± 14.7 *edu*, to 1000 mM, 143.1 ± 7.2 *edu* (p<<0.001). The ratio between the target signal and nuclear background event densities, or the “signal-to-background ratio” (S/B), exhibited the most favorable relationship in *δds* state at 1 M NaCl, 56.5 ± 3.4, Figure 3g. This corresponds to an approximate 5.7x improvement in signal to background event ratio compared to control *ss* in 137 mM NaCl measurements. A slightly lower S/B was achieved with *ss* D3 at 1 M NaCl, 33.1 ± 2.1 (p << 0.001) as a result of the dip in target localizations under those conditions. An ∼2.5-fold improvement was also observed with respect to the ratio between the target signal and cytoplasmic background event densities, Supplementary Figure 5a. Comparison of cytoplasmic to nuclear background levels quantified by the ratio between event densities for (mitochondria-distal) cytoplasmic and nuclear regions exhibited a general trend towards unity when using the *δds* state systems, Supplementary Figure 5b.

To evaluate the generality of these findings we additionally tested a different strand sequence designed for Shielded DNA-PAINT, the D2 docking sequence (see Supplementary Figure 1) attached to secondary ABs in *ss & δds* incubated states at 1 M NaCl, Supplementary Figure 6. An approximate 2.5 fold reduction in the non-specific nuclear event densities occurred when incubating the DNA-PAINT D2 secondary in 1 M NaCl compared to 137 mM NaCl, but without shield strands. In the *δds* incubated state (i.e. with shield strands) at 1 M NaCl the reduction was ∼4.4 fold, Supplementary Figure 6a. Similar to D3 ABs, the cytoplasmic background was reduced ∼2.4x and the target signal was relatively unchanged between control and *δds* state, Supplementary Figure 6b-c. Overall, signal to background ratios for nuclear and cytoplasmic regions saw broadly similar improvements as observed with the D3 docking strand and the ratio between nuclear and (mitochondria distal) cytoplasmic regions got closer to unity with Shielded DNA-PAINT incubation, Supplementary Figure 6d-f. For a full list of measurements see Supplementary Table 2.

Our D2 and D3 sequences are longer (28-30 bp) than typical DNA-PAINT markers (9-11 bp) to enable the ability to functionalize the marker for various applications^7,9,11^ that generally require longer DNA docking strands. For example, the D2 docking strand design was previously used to functionalize markers for repeat docking domains^7^ and also formed one half of the base used to create a super-resolution proximity sensor^11^. To enable the use of these schemes, post Shielded DNA-PAINT labeling, when the D2 or D3 strands are in *δds* state one can safely remove them through the process of toehold-mediated strand displacement^21^ in order to further functionalize the marker (as we have previously demonstrated, for example, with Repeat DNA-PAINT, Fig. 1c in ^11^). For completeness, we also tested antibodies conjugated with a shorter docking strand that is 11 bp long, D3s, and compatible with the D3 imager. These secondary ABs also accumulated within the nucleus, Figure 4a. We adapted Shielded DNA-PAINT to the shorter docking strand, employing a fully complementary sequence to shield the DNA-PAINT marker during its incubation with the sample, Figure 4b, which reduced the nuclear background, Figure 4c. Due to the shorter sequence the shield strand is expected to hybridize only transiently with the shorter D3s strand (melting temperature ∼36 °C at 1 M NaCl) however, when applied in excess creates a system where the D3s docking strands are almost continually occupied. Washing then effectively removes the shield strand in this scenario, enabling imager D3 to access the DNA-PAINT D3s docking strands. Super-resolution measurements of D3s, Figure 4d, in these two configurations reported nuclear event densities of 10.8 ± 0.6 *edu* in *ss* and 2.3 ± 0.3 *edu* when *ds*, a ∼5x reduction, a level similar to that achieved with D2 and D3 docking strands. This equates to an improved S/B relationship of ∼5.8 times, Figure 4e. We did observe a moderate decrease in average photon rates from 3.9k photons/frame with the D3 docking strand versus 3.2k photons/frame using the D3s docking strand (which has docking strand and imager dyes in closer proximity) but note that variations from experiment to experiment were larger than the mean differences observed, Supplementary Figure 2b.

**Figure 4.**
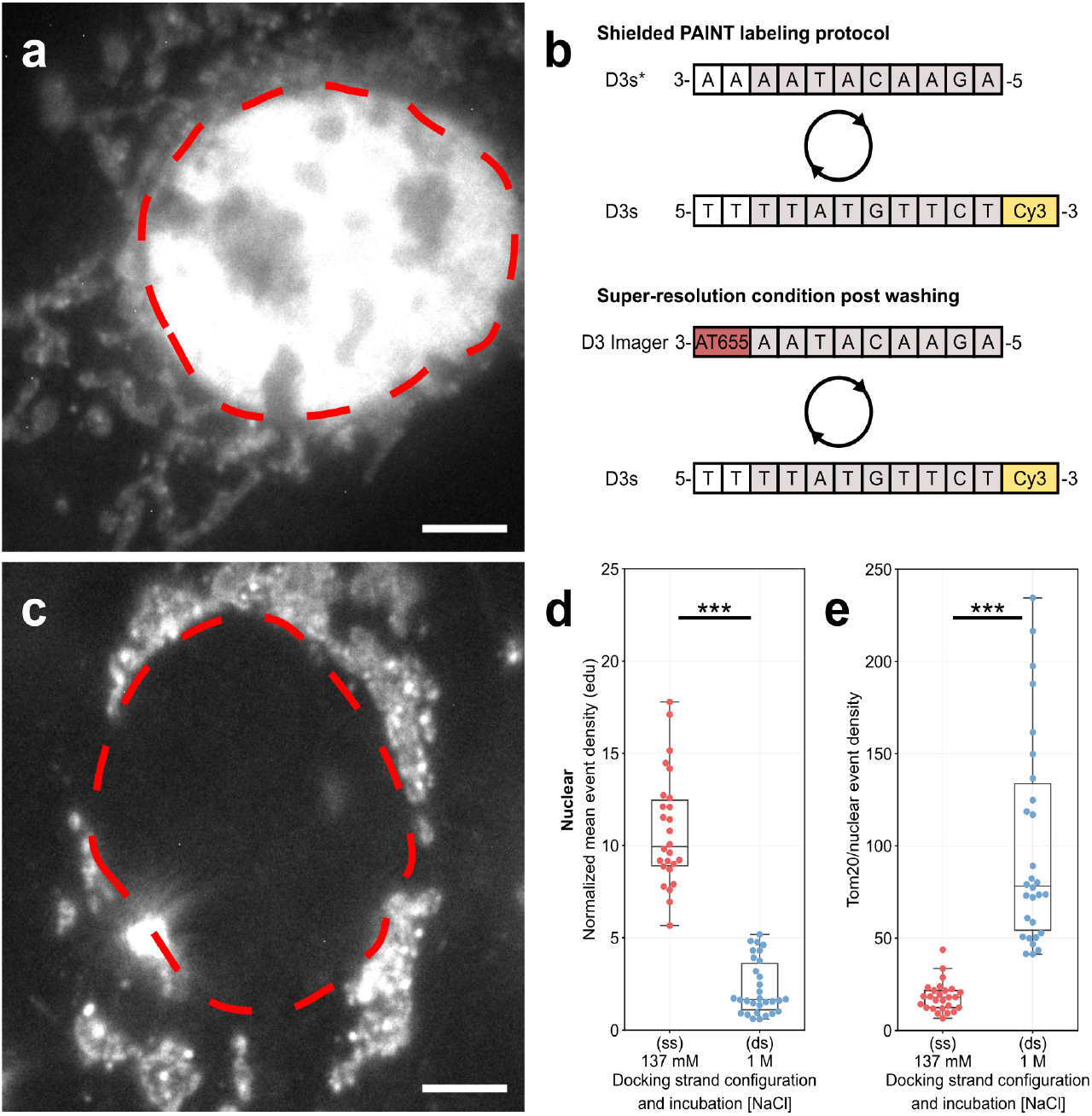
Shielded DNA-PAINT labeling with a short 11 nt, D3s, sequence. **a**. At low 137 mM NaCl, D3s binds to the nucleus. A schematic diagram of the short oligonucleotide sequences, **b**, shows a fully complementary D3s* used to occupy the D3s docking strand whilst labeling the sample. At this size the binding is transient or reversible and easily removed through washing following incubation. This then enables D3 imager to bind to D3s docking strands for DNA-PAINT super-resolution single molecule experiments. Applica<on of Shielded DNA-PAINT protocol with this shorter D3s sequence protects the marker from non-specifically binding to the nucleus, **c**, as observed with widefield images. **d**. Nuclear event densities were reduced ∼5 fold equating to a ∼5 fold improvement in Tom20/nuclear signal, **e**, with *ds* 1 M compared to *ss* 137 mM. Red dashed lines in *a & c* indicate the nuclear perimeter. (26 & 30 cells for *ss* 137 mM and *ds* 1 M NaCl experiments, *n* = 3. Boxplot points represent the mean value obtained per measured cell, and line indicates the median. Independent two-tailed t-tests indicate the following levels of significance: *** p ≤ 0.001). Scale bars: 5 μm.

For comparison with L-DNA based DNA-PAINT markers and imagers we also evaluated secondary ABs conjugated with LB12 docking strands (that were modified with a Cy3 dye, see Table 1) and probed these with LP12 imagers. We found that left-handed oligonucleotides conjugated to secondary antibodies also suffered from non-specific uptake during the labeling period, see Supplementary Figure 7a, at a level of 4.8 *edu*. Consistent with our R-DNA based experiments above, this marker-based background is >10 times higher than the imager-only background from LP12 imagers alone (which was 0.27 *edu*, see Fig. 2i). Comparison with Fig. 2 shows that Shielded DNA-PAINT with 1 M NaCl in the incubation buffer and using the (conventional R-DNA based) D2 and D3 DNA-PAINT antibodies reduced the nuclear background levels to similarly low values (<4.5 *edu*) that are otherwise only achievable with the much more expensive L-DNA.

The LB12 AB associated nuclear event density was reduced further to 1.5 ± 0.2 *edu* by increasing the incubation solution NaCl concentration to 1 M. This was still ∼5x greater than the background levels from LP12 imagers alone, see Figure 2a. Target and cytoplasmic event densities hinted at a slight, but statistically non-significant, reduction at 1 M, Supplementary Figure 7b & c. Conventional labeling achieved a target/nuclear ratio of 39.5 ± 2.6, Supplementary Figure 7d, notably ∼1.4x worse than shielded *δds* D3 AB labeling a 1 M NaCl, Figure 3g. With 1 M NaCl incubation the S/B of L-DNA measurements was further improved ∼2.8 times to ∼100.

We considered the possibility that the docking strand dye modification could be exacerbating the non-specific uptake of our DNA conjugated ABs. To test this hypothesis, we conjugated a blank, not dye modified, D3 sequence to a blank secondary antibody. We visualized where the conjugate ended up by using a tertiary labeling approach with Alexa Fluor 488, Supplementary Figure 8a. Under conventional labeling procedures the Alexa Fluor 488 was strongly absorbed into nuclear regions, Supplementary Figure 8b. By using the Shielded DNA-PAINT labeling method when applying the DNA-PAINT secondary, only the target was fluorescently labeled when the tertiary antibody was added, Supplementary Figure 8c. Nuclear uptake led to event densities for *ss* incubated at 137 mM NaCl at 14.2 ± 2.2 *edu*, which was significantly reduced to 1.1 ± 0.2 *edu* (p << 0.001) for *δds* 1 M, Supplementary Figure 8d. This corresponds to an approximate 12-fold reduction in nuclear non-specific events, a further 2x improvement over the dye modified version of the same strand, approaching levels of imager-only interactions (which always contribute to the measured event densities). Tom20 localizations, Supplementary Figure 8e, were relatively similar between conditions 96.4 ± 9.0 *edu* for *ss* 137 mM NaCl compared to 104.1 ± 10.6 *edu* (p = 0.583) for *δds* 1 M NaCl. These findings are thus broadly similar to the observations with Cy3 modified docking strands, omission of the Cy3 modification moderately reduces backgrounds albeit at the expense of no intrinsic widefield signal prior to super-resolution imaging with DNA-PAINT.

### Imaging of Nucleolin with Shielded DNA-PAINT

To illustrate the utility of the Shielded DNA-PAINT labeling approach we applied it to image nucleolin in fixed COS-7 cells, Figure 5. Labeling of this nuclear target with *ss* D3 in standard 137 mM NaCl demonstrates the difficulty of applying conventional DNA-PAINT labeling procedures to imaging nuclear targets, Figure 5ai. The widefield and super-resolution data, Figure 5aii, exhibit substantial nuclear signal and notably also considerable cytoplasmic signal. DNA-PAINT markers, using the same D3 conjugation, but now incubated in a *δds* state at 1 M NaCl, Figure 5bi-ii, leave the nucleoli within the nucleus clearly visible above a lower level nucleolin signal typically present across the nucleus. Note that this lower level signal across the nucleus corresponds to nucleolin in the nucleus that is expressed outside of the nucleoli and has been observed previously with conventional antibody staining (see, for example. Fig. 1 in Terrier et al^22^). A cytoplasmic background signal surrounding the cell when using conventional staining conditions (Figure 5ai,ii) is no longer discernable in widefield images, Figure 5bi, indicating the effectiveness of the Shielded DNA-PAINT protocol. This is mirrored in the super-resolution image where the nucleoli are well defined and cytoplasmic signal is, as expected, low.

**Figure 5.**
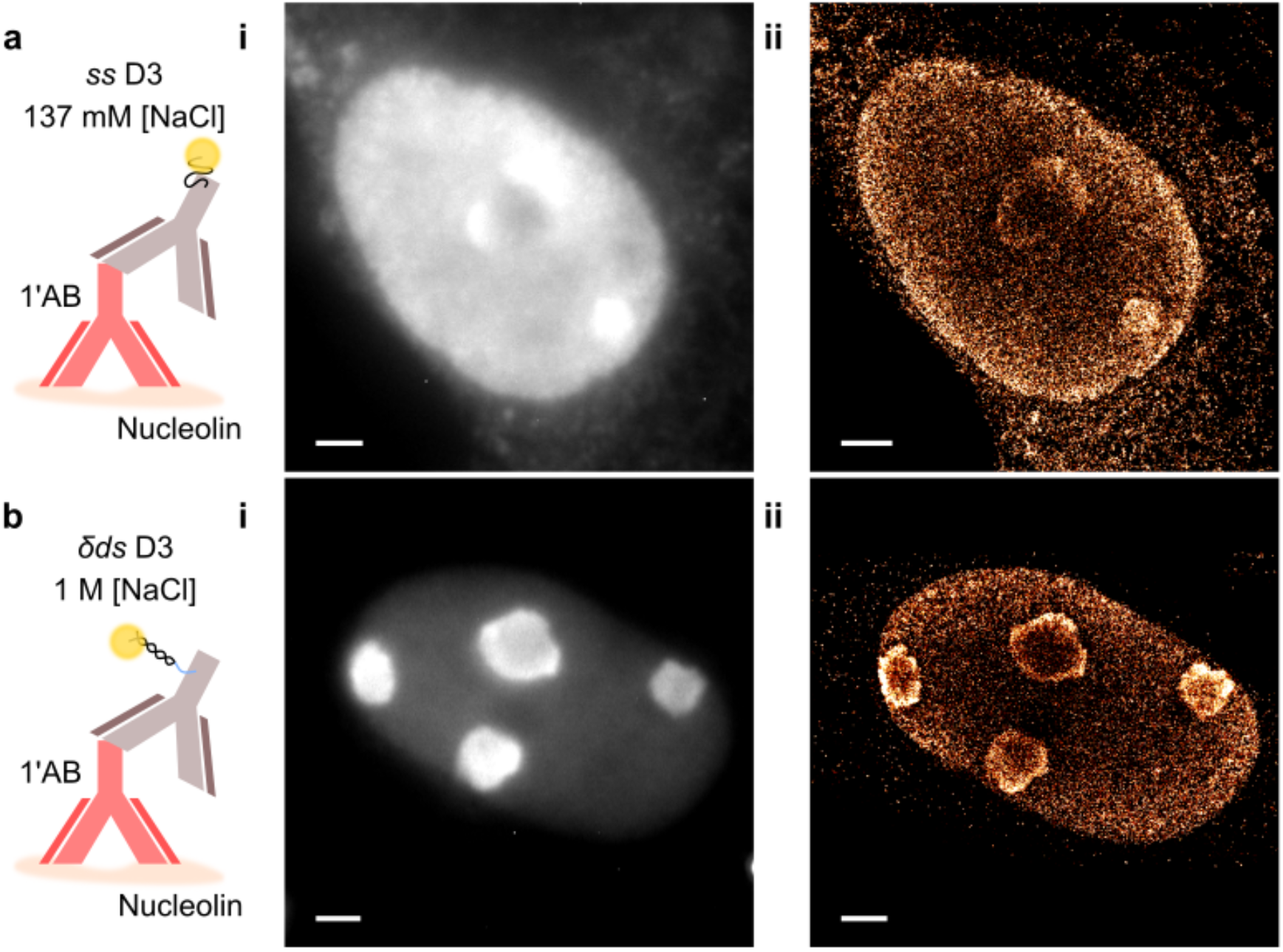
DNA-PAINT labeling of nucleolin staining in COS-7 cells. **a**. Schematic of *ss* D3 incubated in 137 mM NaCl. As expected, the DNA-marker was observed in widefield images, **ai**, to non-specifically incorporate itself across the nucleus of the cell and across the cytoplasm. Super-resolution measurements, **aii**, of the same region match the non-specific signal seen in the widefield image. **b**. Schematic of *δds* D3 incubated in 1 M NaCl incubation solution where the light-blue portion of the *δds* indicates where the imager binds. **bi**. Widefield images show the nucleoli present within the nucleus of cells with much greater contrast. **bii**. Super-resolution detection of single molecule events display well-defined nucleoli and minimal cytoplasmic localizations. Scale bars: 2 μm.

## Discussion

While background in the nucleus of mammalian cells due to non-specific imager binding had been observed during DNA-PAINT imaging before^16^, the much larger contribution from DNA-PAINT AB retention had not been recognized as such. Background from antibodies exhibiting *ss* DNA had been previously noticed in widefield images during multiplexed imaging for many protein targets^23,24^ but had not been quantified in DNA-PAINT imaging. It is highly relevant in practice since we here show that these (marker-based) backgrounds typically are at least ten times higher than imager backgrounds alone. We essentially resolve this problem by introducing a Shielded DNA-PAINT protocol with modified incubation solution that greatly reduces these backgrounds to a low level similar to that achievable with L-DNA probes but using much more widely available conventional right-handed DNA probes.

Key to routinely visualize and easily detect this non-specific background with basic widefield fluorescence imaging was using dye modified DNA-PAINT markers which revealed substantial accumulation of the dye within nuclear regions of cultured cells. DNA-PAINT measurements corroborated the widefield information with event densities about an order of magnitude greater than from imagers only. This also implies that maneuvers aimed at reducing imager concentration through the use of repeated docking domains^7^ have little effect on the dominating marker-based backgrounds as the reduction in imager concentration in this method is compensated by the increase in the number of docking domains per secondary AB.

In general, a sequence dependence of non-specific retention of DNA-PAINT antibodies seems difficult to predict. There are likely differences that depend both on strand length and properties of specific sequences. For example, the degree to which some of the non-specific labeling was reduced, when increasing the NaCl concentration of incubation solutions, differed for the sequences that we evaluated when docking strands remained single-stranded (no shielding), e.g. when comparing D2 and D3 conjugated ABs.

By contrast, Shielded DNA-PAINT labeling, the application of partially or fully complementary oligonucleotides to DNA-PAINT docking strands prior to being added to the sample, consistently reduced non-specific event densities in both nuclear and cytoplasmic regions of cells. Further, with Shielded DNA-PAINT we saw no detectable decrease in specific event densities at target locations (as was seen with 1M NaCl and *ss* docking strands), overall leading to improved signal to background ratios when used with 1 M NaCl in the incubation solution.

Experiments with docking strands lacking a dye modification suggested that the presence of the Cy3 dye was not a major contributor to non-specific retention of the marker. Nevertheless, the improvement in target/nuclear ratios (signal to background) was ∼2x more favorable with a plain docking strand D3 versus D3 with dye modification. Despite this finding, the ability to observe DNA-PAINT markers prior to super-resolution imaging in widefield mode is extremely useful and the dye modified oligos can also assist in the quantification of the fraction of oligos successfully conjugated to the marker of choice^11^.

In agreement with a prior study^16^, an L-DNA imager, LP12 imager, performed the best in terms of low levels of non-specific binding within the nucleus. Consistent with ABs conjugated to conventional right-handed DNA, elevated nuclear event densities were detected following conventional incubation of the L-DNA marker with LB12 docking strands. The marker-based L-DNA background was 16x higher than the LP12 imager-only background. In absolute terms, the background of <5 *edu* is sufficiently low that it will not interfere with moderately expressed target protein signals (S/B is ∼40 for the Tom20 target signal).

The reduction in background achieved with Shielded DNA-PAINT becomes clear by comparison with the LB12 DNA-PAINT AB performance as a reference. Nuclear marker-based backgrounds were reduced broadly from levels of 15-25 *edu* to ∼4 *edu*, smaller than the level seen with LB12 ABs (at standard NaCl). This reduction corresponds to ∼5.7x and ∼2.5x improved target/nuclear and target/cytoplasm ratio, respectively, as measures of desired target signal to backgrounds. The S/B ratios are in a range of 30-50, in some cases >100 when related to the (relatively abundant) Tom20 target. The remaining AB associated background is in a range that is only 2-4 times larger than imager-only backgrounds. In the case of D3 without a dye the remaining background is as small as 1.1 *edu* and likely mostly reflects remaining imager-only backgrounds.

The reduced non-specific, nuclear accumulation of (partially) double-stranded markers compared to classic single-stranded constructs could be due to a variety of factors. Primarily, we hypothesize that hydrophobic or hydrogen-bonding interactions may occur between the unpaired nucleotides of ssDNA markers and either proteins or other nucleic acids in the nucleus, which would not occur, or would be reduced, in dsDNA constructs.

Coulomb interactions may also play a role, for instance with positively charged DNA binding proteins. Regardless of the source of interaction, it is likely that the very different polymeric properties of single-stranded and double-stranded DNA, with the former being highly flexible (persistence length ∼2 nm) and the latter much more rigid (persistence length ∼50 nm), modulates non-specific interactions^25,26^. One may indeed envisage flexible ssDNA being able to fold into configurations maximizing affinity with non-specific targets, which may be inaccessible to dsDNA.

Evidence that increasing ionic strength reduces nuclear accumulation in single-stranded markers supports the hypothesis that Coulomb interactions may be involved, which become screened at high salt concentration. It is however also possible that a higher ionic strength hinders hydrophobic, or hydrogen bonding interactions mediated by the nucleobases by stabilizing secondary structures within the single-stranded docking strand^27^ or interactions between the docking strand and hydrophobic or hydrogen-bonding residues on the secondary antibody.

It is possible that binding of markers non-specifically in the nucleus (or the cytoplasm) may be increased when multiple ssDNA strands are present on a single marker (as may occur with random-labeling) and that this involves cooperative binding in some way. In this scenario the use of site-specific docking strand attachment, as for example regularly employed with nanobodies^28^, may reduce marker-based backgrounds to the level of imager-based backgrounds. Accordingly, such marker systems should be investigated with the methods presented here.

Alternative blocking strategies could be considered, such as the addition of charged polymer dextrans^20^ or salmon sperm DNA^23^. Exploratory experiments with a charged polymer dextran, >500 kD dextran sulfate, using concentrations sufficient to reduce non-specific nuclear labeling, exhibited a concomitant reduction in target labeling so that S/B ratios were substantially smaller than achieved with Shielded DNA-PAINT (Supplementary Figure 9). We therefore did not pursue these approaches further given the comparative simplicity and effectiveness of Shielded DNA-PAINT protocols presented here.

For highest signal to background ratios L-DNA markers with additional charge screening by incubating the DNA-PAINT ABs in 1 M NaCl are still a possible choice. In practice Shielded DNA-PAINT with conventional DNA docking strands should be suitable for observing all but the most sparsely expressed target proteins within the nucleus.

## Conclusion

We have shown that during the labeling procedure with DNA-PAINT ABs there can be significant non-specific retention of these markers when applying them in a conventional antibody incubation solution. By incubating the markers instead in a partially double-stranded (*δds*) or fully double-stranded configuration and increasing ionic strength of the incubation buffer, Shielded DNA-PAINT labeling significantly reduces non-specific uptake of DNA markers into the nucleus and cytoplasm of cells. This approach helps to reduce marker-based backgrounds (DNA-PAINT AB retention) to levels similar to the lesser imager-only contributions. In so doing, our approach can be considered by anyone conducting DNA-PAINT based experiments on biological specimens. Notably, the level of marker-based backgrounds should be evaluated routinely in DNA-PAINT experiments. Shielded DNA-PAINT labeling performed similarly to conventional labeling using L-DNA markers albeit at a fraction of the cost. This approach is straightforward to incorporate into existing super-resolution labeling protocols and enables the use of conventional right-handed DNA. The variable length of docking strands used in this study are compatible with the concept of fluorogenic imager use and would further help to reduce non-specific interactions for Shielded DNA-PAINT labeling applications^9^.

## Methods

### Oligonucleotides

The oligonucleotides used in this study were first checked with NUPACK analysis web application (www.nupack.org) to determine likely folding behaviors and to check complementarity with additional sequences. Docking strands (DS) were ordered from IDT whilst imagers and non-modified strands were ordered from Eurofins. Left-handed DNA (L-DNA) were ordered from Biomers.net with the same modifications as their right-handed counterparts, see Table 1 for a full list of sequences used. Docking strands had a 5’ Azide modification and 3’ fluorophore modification of Cyanine 3. Imagers had a 3’ ATTO 655 modification which when hybridized were detected as single molecule events.

The D2 design was previously used to functionalize markers for repeat docking domains^7^ and also formed one half of the base used to create a super-resolution proximity sensor^11^. The D3 imager was previously introduced as P39* in another study^13^ and we incorporated the corresponding complementary domain into the D3 docking strand.

### Conjugation of Docking Strands to Antibodies

Blank AffiniPure goat anti-mouse (#115-005-003) or anti-rabbit (#111-005-003) IgG (H+L) secondary antibodies (Jackson ImmunoResearch, PA) were conjugated to azide modified docking strands following click-chemistry protocols previously described by Schnitzbauer et al^6^. Briefly, antibodies were labelled with NHS-sulfo-Ester DBCO for 45 minutes at room temperature. 10 minutes incubation with 80 mM Tris (ThermoFisher Scientific, #15568-025) was used to stop the reaction. 7k MWCO Zeba™ spin columns (ThermoFisher Scientific, #89882) were used to remove unbound DBCO. Oligonucleotides were incubated overnight with the DBCO-antibodies at 4°C at a concentration aimed at achieving ∼2 oligos per antibody. A final purification step using Amicon 100 k spin columns (Merck, #UFC510024) was used to remove unbound oligonucleotides. Newly conjugated antibodies were diluted 1 mg/mL in PBS containing 0.05% sodium azide and were subsequently aliquoted and stored at -20°C until use. For dye modified oligonucleotide conjugations the degree of oligo-antibody labeling was determined by measuring the dye absorbance peaks before versus after conjugation and purification, (N60-Touch, Fisher Scientific). We typically achieve 1-3 oligo per antibody, Supplementary Figure 10, see also^11^.

### Cell Culture

COS-7 cells (Cytion, #605470) were grown at 37 °C in a 5% CO_2_ humidified incubator. The cells were maintained in DMEM:Ham’s F12 (Merck, 51445C) supplemented with 10% fetal bovine serum (Merck, #F7524), 100 units/mL penicillin, 100 μg/mL streptomycin, and 0.25 μg/mL amphotericin B. During subculturing, cells were washed with PBS (free of Ca^2+^ and Mg^2+^) and incubated with cell dissociation agent TrypLE (Thermofisher, 12604013) for ∼5 minutes. Cells were seeded on pre-sterilized number 1.5 glass coverslips (Menzel-Gläser 22×22 mm) and grown for 24-48 hours to achieve ∼70% confluency. Once the medium was removed, cells were fixed following a protocol as in Jimenez et al^29^. Briefly, the cells were fixed for 10 minutes in PEM buffer (80 mM PIPES, 5 mM EGTA, 2 mM MgCl_2_) containing 4% formaldehyde (FA) (32% sol. in glass ampoules, Electron Microscopy Sciences, #50980494) with 4% sucrose (Sigma, #S0389). Samples were then washed >3 times in PBS. Coverslips were then mounted onto custom-made 3mm thick Perspex chambers for easy access solution changes.

### Labeling Procedure

Fixed cells were permeabilized and blocked for 10 minutes in phosphate buffered saline, PBS (Sigma, #P4417), containing 0.1% Triton X-100 (Sigma, #T9284) and 2.5% BSA (Merck Millipore, Probumin #81-066-4). Primary antibodies (Santa Cruz Biotechnology, mouse monoclonal IgG_2a_ Tom20 (F-10), #sc-17764 or rabbit polyclonal IgG Tom20 (FL-145), #sc-11415) diluted 1:1000 in incubation solution containing 2% BSA, 2% normal goat serum, 0.05% Triton X-100, 0.05% sodium azide (Sigma, #S2002) were incubated with the sample for 1 H at RT. Samples were then washed >3 times in PBS. For single-stranded (*ss*) experiments the DNA-PAINT secondary antibodies were diluted in incubation solution (1:200) containing either 137 mM, 500 mM or 1 M total NaCl (Sigma, #S7653) concentration. Shielded DNA-PAINT labeling protocol involving partially double-stranded (*δds*) or *ds* experiments were additionally incubated with their corresponding oligonucleotide (∼500 nM) to complement the docking strand on the secondary antibody. These solutions were mixed for 10-20 minutes at RT to accomplish appropriately high hybridization coverage. The samples were incubated with either *ss, δds*, or ds in the specific salt concentration required, depending on experiment, for 1 H at RT. All samples were then washed in imaging buffer, PBS containing 500 mM total NaCl, >3 times over the course of 30 minutes. Secondary only experiments were conducted as above, but with the primary antibody incubation step omitted. Dextran sulfate experiments were conducted with incubation solution containing 137 mM NaCl and diluting dextran sulfate 50% solution (Sigma, #S4031) to either 0.02% or 0.1%. Samples were incubated for 1 H at room temperature and washed as normal.

### Experimental Setup

A modified inverted Nikon Eclipse Ti-E microscope (Nikon, Tokyo) was used to acquire data through a 1.49 NA APO, 60x oil immersion TIRF objective lens (Nikon, Tokyo). Focus was controlled with a piezo objective scanner (P-725, Physik Instrumente, Karlsruhe). Signal was collected with an Zyla 4.2 sCMOS camera (Andor, Belfast). ATTO 655 imagers were excited with a 140 mW LuxX 647 nm CW diode laser (Omikron, Rodgau). Widefield images were achieved using a LED light source (p4000 CoolLED, Andover). An auxiliary camera captured transmitted light at a non-interfering wavelength to correct for focal drift^17,30^. Briefly, a calibration stack of transmitted light images above and below the focal plane were acquired and actively monitored during any super-resolution acquisition. With this system, a 50 nm tolerance of drift in z was actively maintained. Red fluorescent beads ∼200 nm in diameter, (Thermo Scientific, product #F8887), were diluted 1:10k in imaging buffer and added to each sample. The beads were allowed to settle for approximately 15 minutes before excess beads were removed with several light washes with imaging buffer. These fiducial markers were used to correct any lateral drift over the period of an acquisition. Super-resolution data was acquired with a camera integration time of 100 ms. The open-source Python Microscopy Environment (PyME) software (https://github.com/python-microscopy/python-microscopy) and additional functional plug-ins (https://github.com/csoeller/PYME-extra), were used to both acquire and analyze data. Briefly, for super-resolution acquisitions the default parameter settings for DNA-PAINT were used, these included a ‘Point Finding Threshold’ value of 2.0 and ‘Background’ setting of 0:0 (i.e. no sliding background average is subtracted). Single molecule events were fitted using a 2D Gaussian PSF model^31^ (LatGaussFitFR module).

### Data Acquisition

#### Event Density Experiments

Stock imagers were kept frozen at 100 μM, an intermediate working stock imager solution of 100 nM in imaging buffer was used to freshly dilute imagers to 0.1 nM prior to beginning the imaging experiment. Larger volumes were used to reduce pipetting error, nominally 6-10 μl of 100 nM imager were diluted into 6-10 mL total volume imaging buffer. Samples were always imaged with freshly diluted 0.1 nM D2 imager, D3 imager, LP12 imager depending on the experiment. Each acquisition captured at least 5 k frames, up to 25 k, at 100 ms integration time.

#### Non-specific Imager Binding

COS-7 cells were fixed, permeabilized, and blocked as described in the ‘Labeling procedure’. These unlabeled samples were then introduced to 0.1 nM of the specific imager being tested (P1 imager, D2 imager, D3 imager or LP12 imager) in imaging buffer. Five thousand frames were captured at 100 ms frame integration time for each acquisition. Images of non-specific binding for Fig. 2 were Gaussian rendered^32^ with 20 nm pixel size. Example super-resolution images of Tom20 were rendered with 10 nm pixel Jittered Triangulation.

#### Nucleolin Imaging with DNA-PAINT

COS-7 cells were fixed, permeabilized, and blocked as described in the ‘Labeling procedure’. Rabbit polyclonal antibody to nucleolin (Abcam, #ab22758) was incubated for 1H at RT and washed in PBS. Samples were then either incubated in control conditions using antibody incubation solution containing 137 mM NaCl or following the Shielded DNA-PAINT labeling protocol for D3 docking strand using D3 shield (allowing D3 imager access) in 1 M NaCl. Samples were washed with imaging buffer a minimum of three times before introducing the ∼200 nm red fiducial markers. Freshly diluted, 0.05 nM, D3 imager was introduced to the sample. Widefield images of the regions imaged were acquired prior to commencing DNA-PAINT super-resolution measurements using a camera integration time of 500 ms. Nucleolin super-resolution images were Gaussian rendered^32^ with 10 nm pixel size.

### Data Analysis

#### Event Density Measurements

Localization events were drift corrected in PyME by using the fiducial markers. Events from the same detection window, i.e. from a single binding event, were coalesced into a single localization. Single molecule events were then rendered as 20 nm pixel size histograms^32^ and saved as .tif file format. The images were imported into Fiji, ImageJ^33^ using Bioformats^34^. The rectangle selection tool was used to draw a single region of interest over the nuclear area for each dataset. Approximately ten areas were selected for cytoplasmic regions. Experiments labelled with primary antibodies had approximately ten regions covering Tom20 signal traced around the outlines of mitochondria using the polygon selection tool, see also Supplementary Figure 3. The inbuilt ‘ROI Manager’ tool was used to record the location where each selection was made and saved as .zip file for future reference. Each label was then measured and set parameters were recorded, including area and mean gray values. These measurements were stored in .csv file format. Custom Jupyter (https://jupyter.org) notebooks written in python code were used to create plots and further analyze the data to quantify event densities. Tom20 and cytoplasmic measurements were averaged per cell and this value was used in the respective boxplots. Event densities are specified in units of *edu* (event density unit) as detected events per μm^-2^ per 1000 camera frames, with each frame lasting 100 ms, i.e. 1 *edu* = 0.01 events μm^-2^ s^-1^. The area and duration normalization takes into account that in DNA-PAINT the number of events are proportional to the area sampled and the duration of imaging. See also data availability statement.

#### Imager Kinetics

The bright times, τ_b_, were determined using the PyME filter for ‘clumpSize’ on coalesced localization data. This value returns the number of frames an event is recorded over and provides an estimate for how long the imager remained bound to the docking strand. The dissociation rate, k_OFF_ = 1/ τ_b_, could then be calculated using a camera integration time of 0.1s. Photon numbers were acquired by filtering events lasting 3 or more frames. The data from the first and last frames were dropped to account for the imager binding or unbinding from the docking strand. From all remaining detections, the median value was taken for each dataset measured.

### Statistical Analysis

All event densities are given as the mean ± standard error of the mean. Boxplot lines indicate the median. Figure 2i, imager-only experiments for P1, D2, D3 & LP12 imagers measured 20, 20, 24 & 16 cells respectively over 3 independent replicates (abbreviated as n=3 in figure legends). Figure 2iv, secondary only experiments were performed on 27 cells over 3 independent replicates. Supplementary Figure 2a, *ss* vs *δds* for D3 (24 datasets for both configurations, n =3) and for D2 (18 & 12 datasets for *ss* and *δds* respectively, n = 3). Supplementary Figure 2b, photon numbers for D3 and D3s 1 M shielded incubation solutions measured 24 and 30 cells, over 3 independent replicates. Figure 3iv-vii & Supplementary Figure 5, D3 experiments were performed on 28, 22, 27, 28, 27 & 24 cells for conditions *ss* 137 mM, *δds* 137 mM, *ss* 500 mM, *δds* 500 mM, *ss* 1 M & *δds* 1 M respectively over 3 independent replicates. Supplementary Figure 6, D2 experiments were performed on 15, 18 & 12 cells for conditions *ss* 137 mM, *ss* 1 M & *δds* 1 M respectively over 3 independent replicates. Supplementary Figure 7, L-DNA 137 mM experiments were performed on 11 cells, and 1 M incubation on 10 cells over 3 independent replicates. Supplementary Figure 8, no dye D3 *ss* 137 mM experiments were conducted on 10 cells, *δds* 1 M had 9 cells, over 3 independent replicates. Figure 4, D3s *ss* 137 mM measured 26 cells and *ds* 1 M had 30 cells, over 3 independent replicates. Supplementary Figure 9, dextran sulfate with D3 Cy3 experiments were performed on 21 and 17 cells for dextran sulfate concentration 0.02% and 0.1%, n = 2. Independent two-tailed t-tests were used to calculate *p*-values to check for statistical significance using the Python library package SciPy and the ‘stats’ module. A *p*-value > 0.05 was considered to be non-significant, see data availability statement.

## Acknowledgements

This work was supported by Biotechnology and Biological Sciences Research Council (BBSRC), BB/T007176/1. E.L. was supported by a doctoral scholarship from Bio-Techne and the University of Exeter.

## Author contributions

Conceptualization: C.S, A.H.C. Data curation: C.S, A.H.C. Formal analysis: E.L, A.F.E.B, A.H.C. Funding acquisition: S.W, L.M, C.S. Investigation: E.L, A.F.E.B, A.S, A.M, C.E, A.H.C. Methodology: E.L, C.S, A.H.C. Project administration: C.S, A.H.C. Resources: L.M, A.S, A.F.E.B, A.M, E.L, C.S, A.H.C. Software: C.S, A.H.C. Supervision: C.E, S.W, L.M, C.S, A.H.C. Validation: E.L, A.F.E.B, A.M, A.H.C. Visualization: E.L, A.F.E.B, A.S, C.S, A.H.C. Writing: All authors contributed to the writing of the manuscript.

