## Supplemental figures and tables for "Reduced Non-Specific Binding of Super-Resolution DNA-PAINT Markers by Shielded DNA-PAINT Labeling Protocols"

Supplementary figures and tables

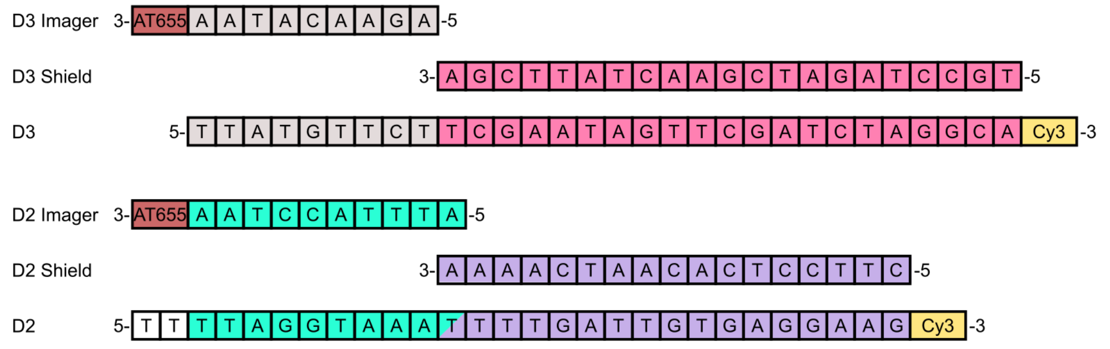


**Supplementary Figure 1.** Schematic diagram of oligonucleotide sequences with binding domains color coded. D3 and D2 imagers bind to the D3 and D2 docking strand portions, respectively, which remain single-stranded when D3 and D2 strands, respectively, are in a δds configuration with their corresponding 'Shield' sequence.

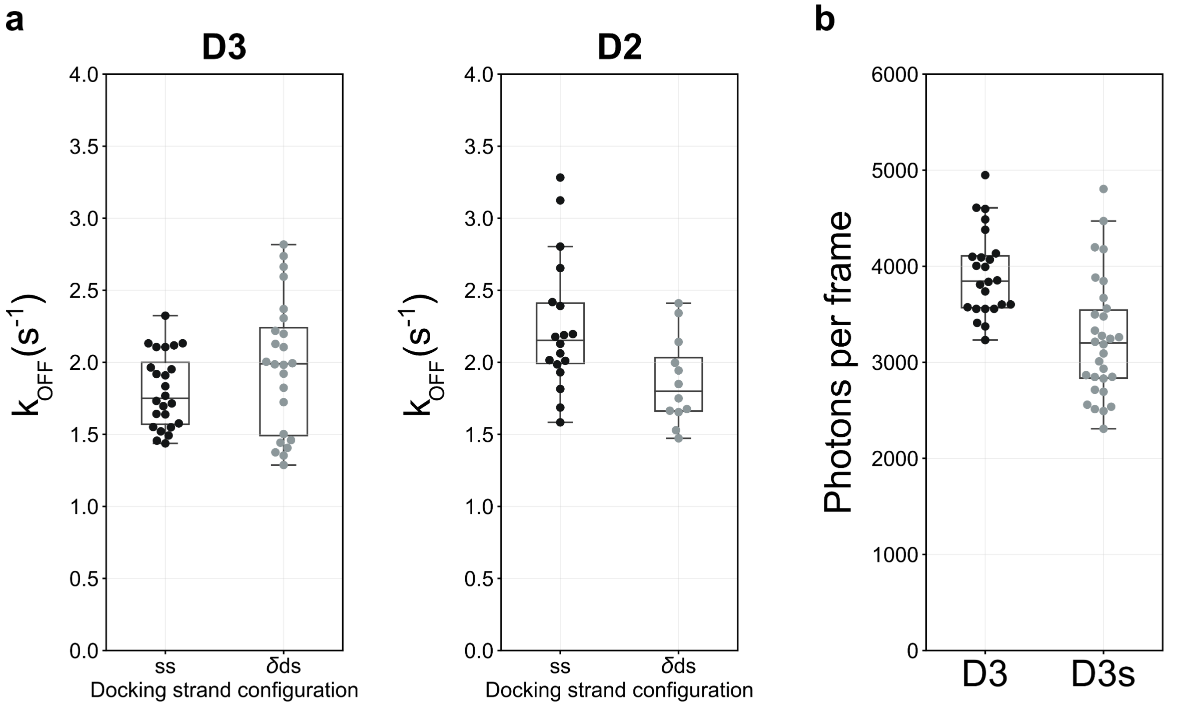


**Supplementary Figure 2. a.** Dissociation rates, $k_{OFF}$, between fully *ss* and *δds* configurations for D3 (mean value ~1.8 $s^{-1}$ and 2.0 $s^{-1}$) and for D2 (mean value ~2.3 $s^{-1}$ and 1.9 $s^{-1}$) exhibit similar rates. The similarity between docking strand configurations indicates that the binding kinetics for the imager is largely unaffected by the '*shielded*' *δds* sequence. Boxplot points represent mean $k_{OFF}$ values calculated per dataset. D3 measurements were taken from 24 datasets for both *ss* and *δds*, n = 3 and for D2, the measurements were taken from 18 and 12 datasets for *ss* and *δds* respectively, n =3. **b.** Detected number of photons per frame for ATTO 655 imager binding to D3 and D3s sequence having a Cy3 dye-modification. D3 mean value 3.9 ± 0.1 k photons compared to D3s mean 3.2 ± 0.1 k (24 vs 30 cells, n =3). Boxplot points represent median number of photons per frame obtained for each dataset measured.

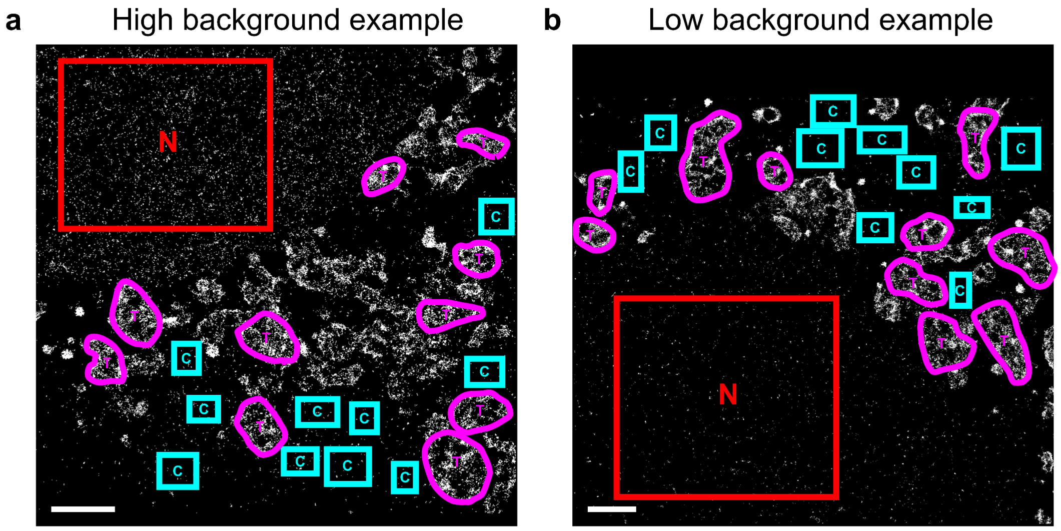


**Supplementary Figure 3.** Super-resolution single molecule event localization images rendered with the histogram method for event density analysis. An example image for both **a.** "high" background, the result of incubating D3 ABs in a *ss* configuration at 500 mM NaCl, and **b.** "low" background, after applying Shielded DNA-PAINT labeling protocols with D3 shield strand incubated at the same 500 mM NaCl concentration in the incubation buffer. In both images, highlighted regions show where measurements were recorded using the Fiji ROI Manager for nuclear (red, 'N'), target signal (magenta, 'T'), and cytosolic (cyan, 'C') areas. Scale bars: 2 µm.

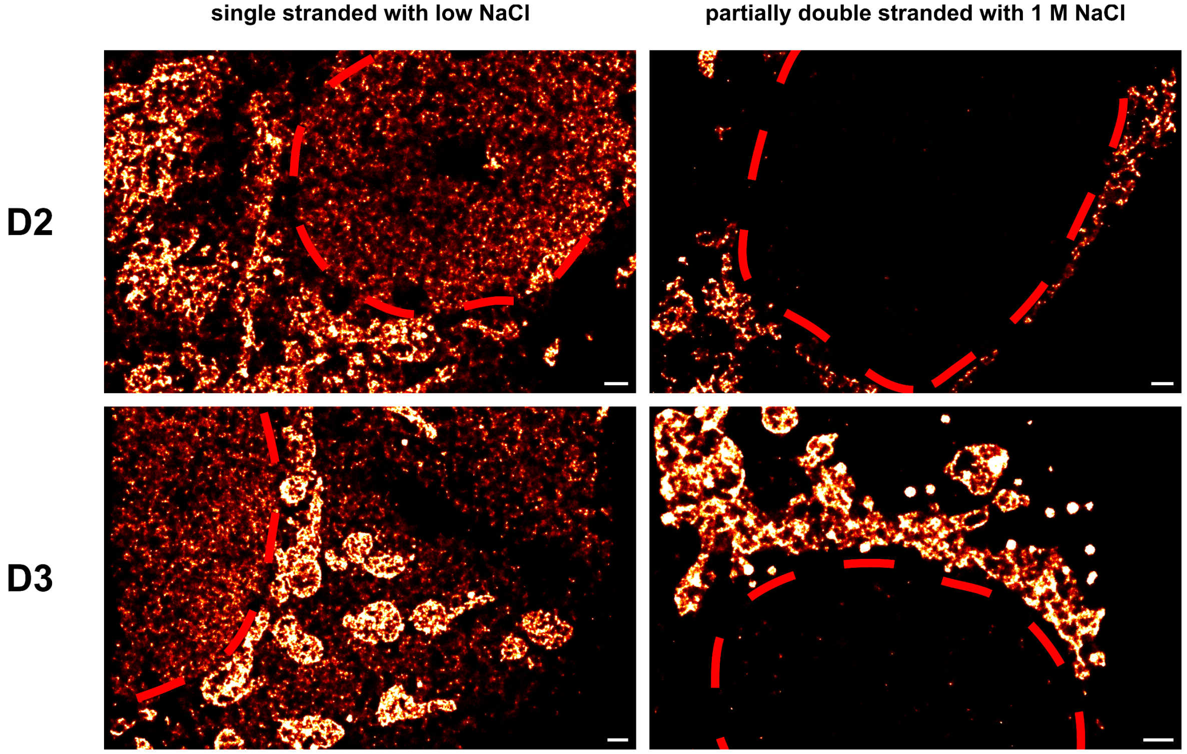


**Supplementary Figure 4.** Super-resolution single molecule event localization images rendered with the jittered triangulation method. The rendered images of example datasets labeled with a primary AB against Tom20 highlight the significant contribution caused by the non-specific attachment of oligo conjugated secondary antibodies at low NaCl (PBS only) incubation solution in single stranded state when compared to the same strands incubated with 1 M total NaCl in a partially double stranded composition. Images share the same intensity look-up-table values. Red-dashed rings indicate the nuclear region of the imaged cell. Scale bars: 1 µm.

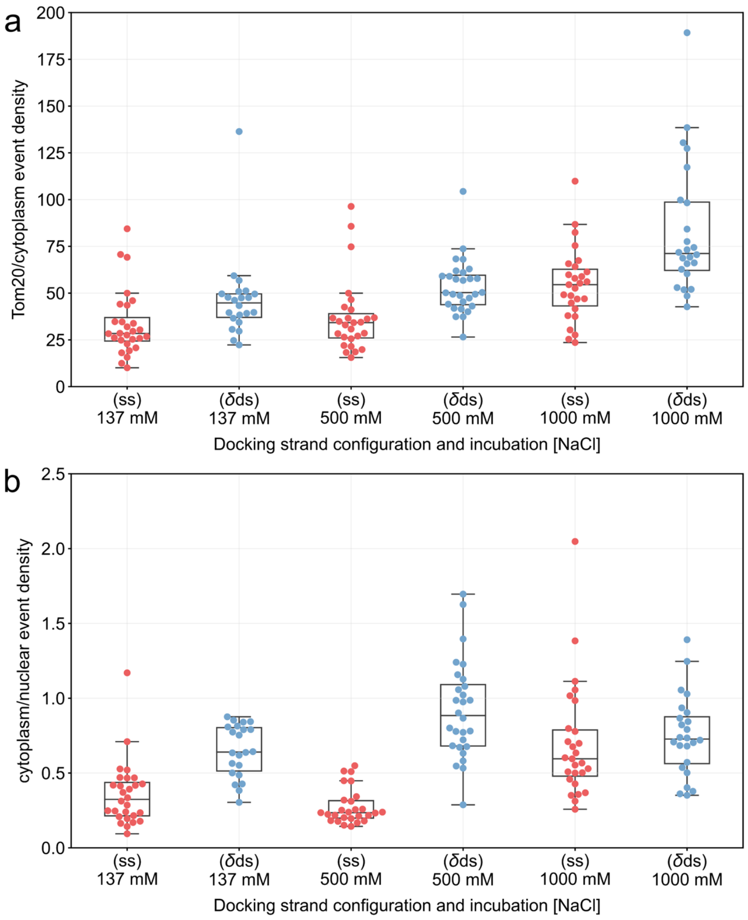


**Supplementary Figure 5.** D3 super-resolution event density ratios for various DNA-marker labeling incubation solutions. **a.** Tom20/cytoplasmic event density ratios exhibit a general increase as NaCl concentration increases from 137 mM to 1 M with *δds* performing better than *ss* configurations. **b**. The ratio of background (cytoplasmic/nuclear) event density contributions display a tendency towards unity as both the nuclear and cytosolic contributions are reduced with interventions during labelling. This shows that the nuclear background is generally reduced to about the remaining cytosolic background level which itself is typically also reduced in magnitude. Accordingly, imaging contrast in the nucleus becomes comparable to imaging contrast in the cytosol using the improved incubation protocols. Boxplot points represent the mean value obtained per measured cell. 28, 22, 27, 28, 27 & 24 cells for conditions as shown left to right, *n* = 3.

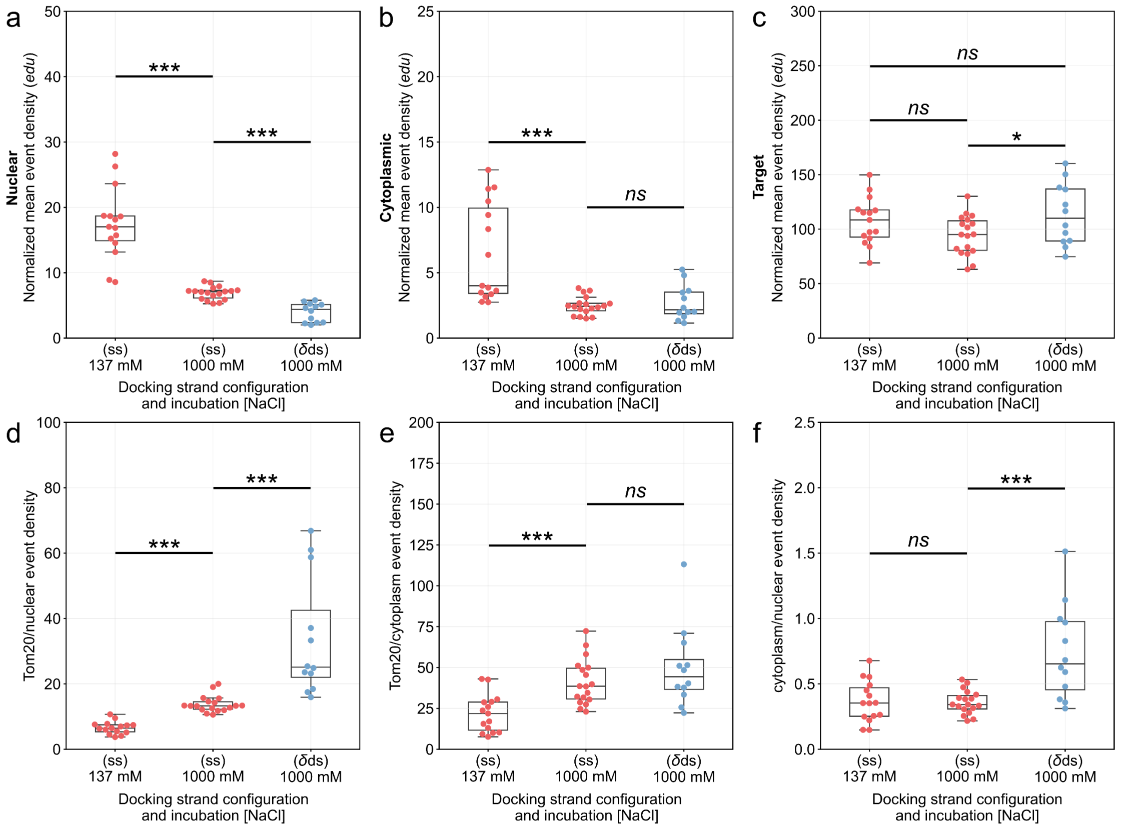


**Supplementary Figure 6.** D2 conjugated AB super-resolution event density measurements for *ss* at 137 mM NaCl, *ss* at 1 M NaCl, and *δds* 1 M NaCl incubation solutions. **a**. Non-specific nuclear event densities decrease with higher NaCl concentration. A *δds* labeling system exhibits the lowest levels of non-specific detection. **b**. Cytoplasmic event densities follow the same behavior, whilst specific Tom20 signal, **c**, maintains comparable levels in each labeling condition. **d**. The ratio of Tom20/nuclear event densities are improved ~5 fold and, **e**, Tom20/cytoplasmic values ~2 fold, similar to D3, between *ss* 137 mM and *δds* 1 M NaCl incubations. **f**. The cytoplasmic/nuclear ratio relationship are unchanged between *ss* at 137 mM and 1 M NaCl, in *δds* configuration at 1 M NaCl non-specific nuclear and cytoplasmic signals become similar to one another. Boxplot points represent the mean value obtained per measured cell. (15, 18, & 10 cells for *ss* 137 mM, ss 1000 mM & *δds* 1000 mM NaCl experiments, *n* = 3; independent two-tailed t-tests indicate the following levels of significance: *ns* $p > 0.05$, * $p \leq 0.05$ and *** $p \leq 0.001$).

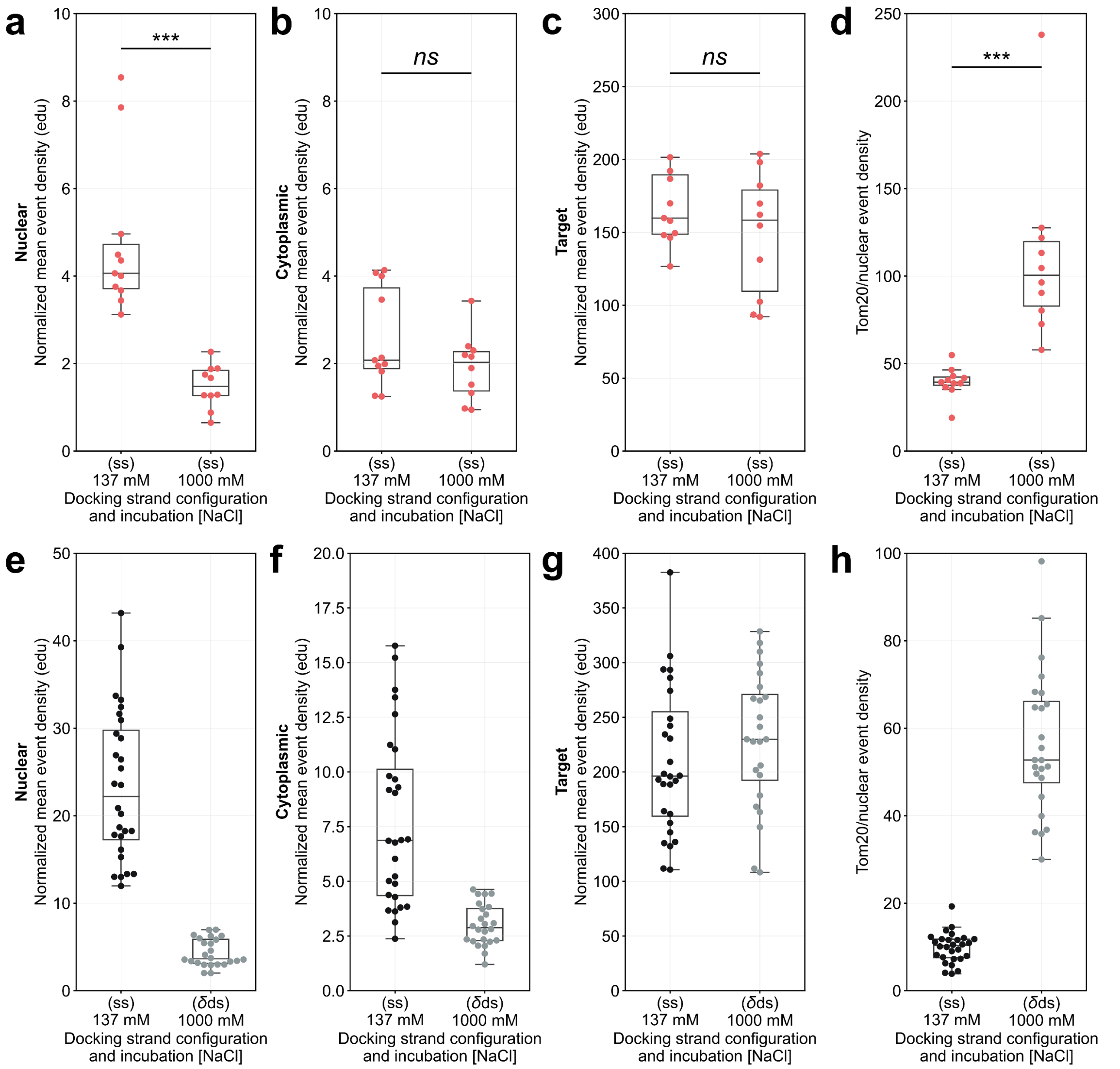


**Supplementary Figure 7**. L-DNA, LB12 modified AB, super-resolution event density measurements for *ss* 137 mM and 1 M NaCl incubation solutions. **a**. Non-specific nuclear event densities decreased ~2 fold at high 1 M NaCl. **b**. Cytoplasmic and **c.** Target Tom20 signals exhibited similar levels at both NaCl concentrations. **d**. Accordingly, Tom20/nuclear event ratios were improved >2 fold when incubated at 1 M NaCl. **e-h.** The equivalent measurements, to the panels directly above, conducted for D3 with docking strands incubated in *ss* 137 mM and *δds* 1 M NaCl, replotted from the main text Figure 3d-g. Boxplot points represent the mean value obtained per measured cell. (L-DNA experiments, 11 & 10 cells for 137 mM and 1000 mM NaCl, *n* = 3; independent two-tailed t-tests indicate the following levels of significance: *ns* p > 0.05 and *** p ≤ 0.001).

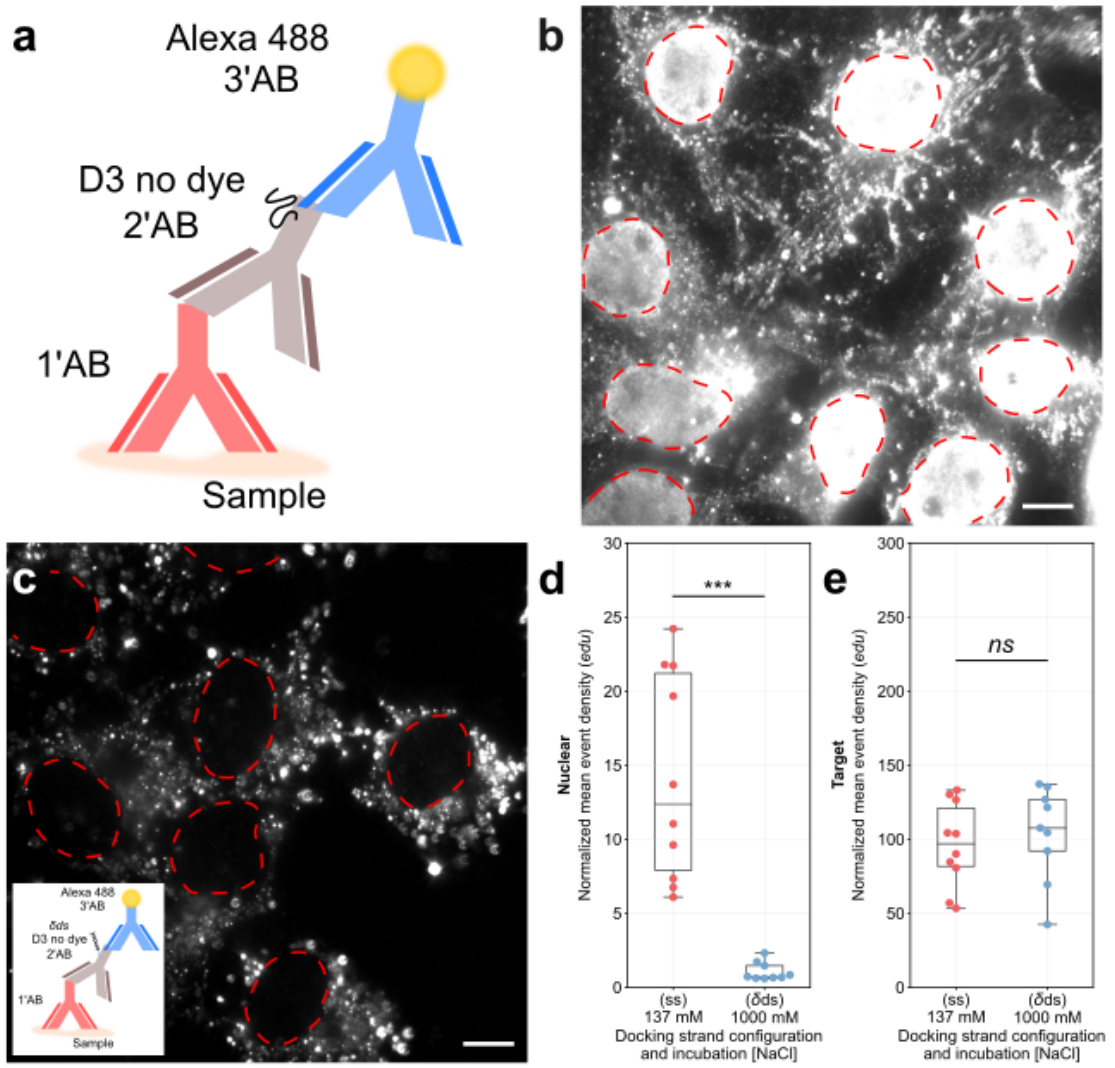


**Supplementary Figure 8.** **"**No dye" DNA-PAINT marker experiments. **a.** Schematic labeling example showing a primary antibody bound to the sample. To this an unmodified D3 oligo (no dye) conjugated to a secondary antibody identifies the primary marker, a commercial Alexa Fluor 488 tertiary antibody shows in widefield images where the secondary antibody binds. **b**. Widefield image of single-stranded, D3 "no dye" secondary ABs bound in the nucleus of COS-7 cells. **c**. Incubation of the D3 "no dye" secondary ABs with D3 docking strands in *δds* state no longer shows a nuclear signal. Inset: Schematic diagram showing the *δds* labeling scenario. **d**. Super-resolution event densities are reduced ~12 fold upon incubation in the *δds* state (Shielded DNA-PAINT at 1 M NaCl) to a level comparable to D3 imager only measurements, suggesting that remaining secondary AB binding in the nucleus is virtually completely abolished. **e**. The target Tom20 signal remains unchanged between conditions. Red dashed lines in *b* & *c* indicate the nuclear perimeter. Boxplot points represent the mean value obtained per measured cell. (10 & 9 cells for *ss* 137 mM and *δds* 1000 mM NaCl experiments, $n = 3$; independent two-tailed t-tests indicate the following levels of significance: *ns* $p > 0.05$ and *** $p \leq 0.001$). Scale bars: 10 µm.

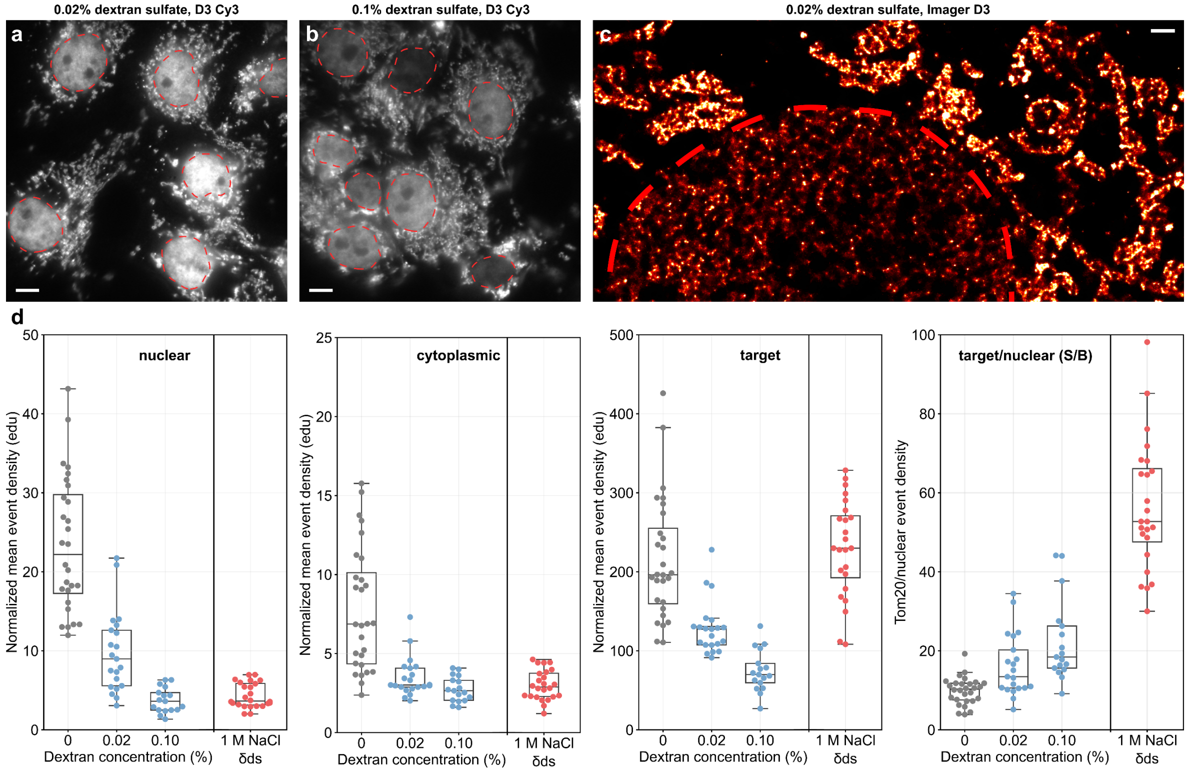


**Supplementary Figure 9.** Experiments using dextran sulfate in secondary antibody incubation solution. **a & b** show widefield images of Tom20 labeled with D3 Cy3 DNA-PAINT secondary using either 0.02% or 0.1% dextran sulfate in the incubation solution during labeling. A slight reduction in signal intensity can be observed at the higher concentration. Widefield images have matching display intensities. **c.** An example super-resolution rendered image of D3 Cy3, incubated in 0.02% dextran sulfate during labeling, highlights that non-specific nuclear interactions are still present. **d.** Measured event densities show an overall decrease in the number of events recorded compared to having no dextran sulfate present, (gray points replotted from the main text Figure 3d-g). Event densities decrease further at the higher, 0.1%, dextran sulfate concentrations. The reduction in non-specific nuclear event densities is offset by the reduction in specific Tom20 target locations leading to minimal improvement in S/B ratios, ~16-22 for 0.02% and 0.1% respectively. For convenience, D3 Cy3 incubated with 1 M NaCl and in a *δds* state has been added to the plots (red points replotted from the main text Figure 3d-g). Boxplot points represent the mean value obtained per measured cell. (21 & 17 cells, n =2). Scale bars: a-b 5 µm, c 1 µm.

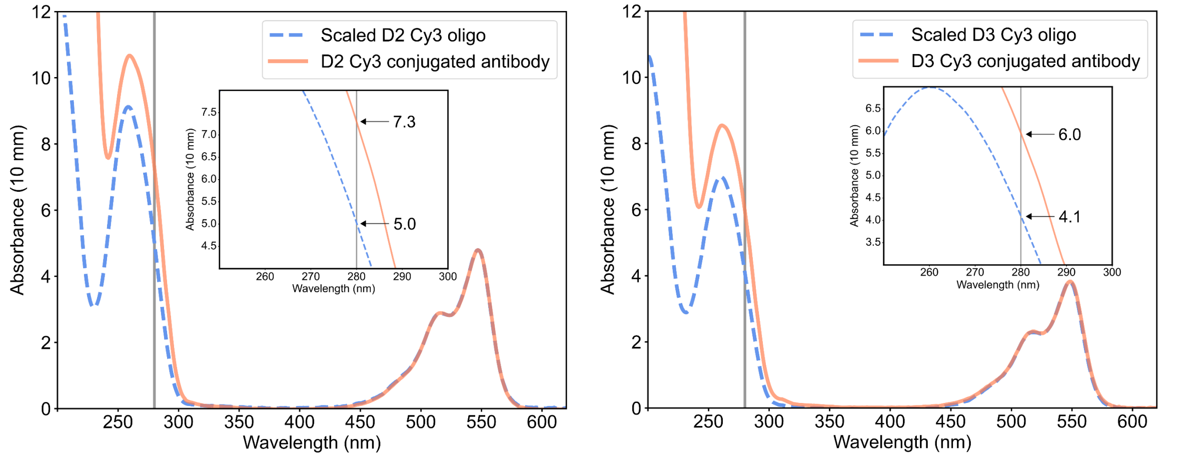


**Supplementary Figure 10.** Absorbance spectra for D2 Cy3 (**left**) and D3 Cy3 (**right**) secondary antibody conjugations. The measurement of the oligonucleotide pre-conjugation is scaled to the final measurement of the completed conjugation using the dye peak from the Cy3 modification. The difference then, between the spectra at the 280 nm value (see inset), provides an estimate for the antibody concentration. In conjuction with the estimate of the dye concentration, using the dye peak, an estimate for how many oligonucleotides present per antibody can be calculated. In this example, we achieved ~2.7 and ~2.8 oligos per antibody for D2 Cy3 and D3 Cy3 conjugations respectively. This approach assumes that each bound oligonucleotide has a dye modification.

| Imager strand | Nuclear mean event density (events µm$^{-2}$ (100 s)$^{-1}$) |
|---|---|
| P1 imager | 2.30 ± 0.13 |
| D2 imager | 1.58 ± 0.17 |
| D3 imager | 0.93 ± 0.08 |
| LP12 imager | 0.27 ± 0.02 |

**Supplementary Table 1:** Numerical data for imager only experiments. All values given as mean ± sem.

| Strand | Incubated state | Mean event density (events µm$^{-2}$ (100 s)$^{-1}$) | | | Mean event density ratios | | |
|---|---|---|---|---|---|---|---|
| | | Nuclear | Cytoplasm | Target | Target/Nuclear | Target/Cytoplasm | Cytoplasm/Nuclear |
| D3 Cy3 | (*ss*) 137 mM | 23.4 ± 1.6 | 7.8 ± 0.7 | 215.4 ± 14.7 | 9.9 ± 0.7 | 33.4 ± 3.3 | 0.36 ± 0.04 |
| | (*δds*) 137 mM | 7.9 ± 0.5 | 5.0 ± 0.3 | 214.5 ± 13.3 | 27.6 ± 1.3 | 46.4 ± 4.8 | 0.65 ± 0.04 |
| | (*ss*) 500 mM | 22.6 ± 1.3 | 6.0 ± 0.5 | 194.2 ± 12.6 | 9.6 ± 0.9 | 37.1 ± 3.8 | 0.27 ± 0.02 |
| | (*δds*) 500 mM | 4.9 ± 0.3 | 4.1 ± 0.2 | 216.1 ± 12.4 | 47.4 ± 3.0 | 53.6 ± 2.8 | 0.92 ± 0.06 |
| | (*ss*) 1 M | 4.7 ± 0.3 | 2.9 ± 0.2 | 143.1 ± 7.2 | 33.1 ± 2.1 | 54.4 ± 3.8 | 0.70 ± 0.07 |
| | (*δds*) 1 M | 4.3 ± 0.3 | 3.0 ± 0.2 | 229.8 ± 12.6 | 56.5 ± 3.4 | 83.1 ± 7.2 | 0.75 ± 0.05 |
| D2 Cy3 | (*ss*) 137 mM | 17.5 ± 1.4 | 6.5 ± 1.0 | 107.4 ± 5.6 | 6.6 ± 0.5 | 21.9 ± 3.0 | 0.37 ± 0.04 |
| | (*ss*) 1 M | 6.9 ± 0.2 | 2.5 ± 0.2 | 94.5 ± 4.3 | 13.8 ± 0.6 | 41.0 ± 3.3 | 0.36 ± 0.02 |
| | (*δds*) 1 M | 4.0 ± 0.4 | 2.7 ± 0.4 | 113.4 ± 8.2 | 33.8 ± 5.3 | 49.8 ± 7.1 | 0.74 ± 0.10 |
| D3 (no dye) | (*ss*) 137 mM | 14.2 ± 2.2 | 1.5 ± 0.1 | 96.4 ± 9.0 | 8.2 ± 1.4 | 69.1 ± 10.7 | 0.13 ± 0.02 |
| | (*δds*) 1 M | 1.1 ± 0.2 | 0.6 ± 0.1 | 104.1 ± 10.6 | 127.4 ± 23.1 | 186.2 ± 28.8 | 0.71 ± 0.09 |
| LB12 Cy3 | (*ss*) 137 mM | 4.8 ± 0.5 | 2.6 ± 0.3 | 179.1 ± 16.6 | 39.5 ± 2.6 | 80.6 ± 10.7 | 0.60 ± 0.09 |
| | (*ss*) 1 M | 1.5 ± 0.2 | 1.9 ± 0.2 | 149.0 ± 13.3 | 110.3 ± 15.8 | 87.5 ± 12.2 | 1.33 ± 0.12 |
| D3s Cy3 | (*ss*) 137 mM | 10.8 ± 0.6 | 2.0 ± 0.1 | 178.9 ± 7.8 | 18.3 ± 1.6 | 98.1 ± 7.4 | 0.20 ± 0.02 |
| | (*ds*) 1 M | 2.3 ± 0.3 | 1.6 ± 0.2 | 170.9 ± 8.3 | 105.6 ± 12.7 | 136.2 ± 12.5 | 0.85 ± 0.14 |

**Supplementary Table 2:** Numerical data for docking strands tested with various incubated states (*ss* vs *δds* or *ds*) and NaCl concentrations. All values given as mean ± sem.
